# Toward an integrated classification of neuronal cell types: morphoelectric and transcriptomic characterization of individual GABAergic cortical neurons

**DOI:** 10.1101/2020.02.03.932244

**Authors:** Nathan W. Gouwens, Staci A. Sorensen, Fahimeh Baftizadeh, Agata Budzillo, Brian R. Lee, Tim Jarsky, Lauren Alfiler, Anton Arkhipov, Katherine Baker, Eliza Barkan, Kyla Berry, Darren Bertagnolli, Kris Bickley, Jasmine Bomben, Thomas Braun, Krissy Brouner, Tamara Casper, Kirsten Crichton, Tanya L. Daigle, Rachel Dalley, Rebecca de Frates, Nick Dee, Tsega Desta, Samuel Dingman Lee, Nadezhda Dotson, Tom Egdorf, Lauren Ellingwood, Rachel Enstrom, Luke Esposito, Colin Farrell, David Feng, Olivia Fong, Rohan Gala, Clare Gamlin, Amanda Gary, Alexandra Glandon, Jeff Goldy, Melissa Gorham, Lucas Graybuck, Hong Gu, Kristen Hadley, Michael J. Hawrylycz, Alex M. Henry, DiJon Hill, Madie Hupp, Sara Kebede, Tae Kyung Kim, Lisa Kim, Matthew Kroll, Changkyu Lee, Katherine E. Link, Matthew Mallory, Rusty Mann, Michelle Maxwell, Medea McGraw, Delissa McMillen, Alice Mukora, Lindsay Ng, Lydia Ng, Kiet Ngo, Philip R. Nicovich, Aaron Oldre, Daniel Park, Hanchuan Peng, Osnat Penn, Thanh Pham, Alice Pom, Lydia Potekhina, Ramkumar Rajanbabu, Shea Ransford, David Reid, Christine Rimorin, Miranda Robertson, Kara Ronellenfitch, Augustin Ruiz, David Sandman, Kimberly Smith, Josef Sulc, Susan M. Sunkin, Aaron Szafer, Michael Tieu, Amy Torkelson, Jessica Trinh, Herman Tung, Wayne Wakeman, Katelyn Ward, Grace Williams, Zhi Zhou, Jonathan Ting, Uygar Sumbul, Ed Lein, Christof Koch, Zizhen Yao, Bosiljka Tasic, Jim Berg, Gabe J. Murphy, Hongkui Zeng

## Abstract

Neurons are frequently classified into distinct groups or cell types on the basis of structural, physiological, or genetic attributes. To better constrain the definition of neuronal cell types, we characterized the transcriptomes and intrinsic physiological properties of over 3,700 GABAergic mouse visual cortical neurons and reconstructed the local morphologies of 350 of those neurons. We found that most transcriptomic types (t-types) occupy specific laminar positions within mouse visual cortex, and many of those t-types exhibit consistent electrophysiological and morphological features. We observed that these properties could vary continuously between t-types, which limited the ability to predict specific t-types from other data modalities. Despite that, the data support the presence of at least 20 interneuron met-types that have congruent morphological, electrophysiological, and transcriptomic properties.

**Highlights:**

- Patch-seq data obtained from *>*3,700 GABAergic cortical interneurons
- Comprehensive characterization of morpho-electric features of transcriptomic types
- 20 interneuron met-types that have congruent properties across data modalities
- Different Sst met-types preferentially innervate different cortical layers

## Introduction

The mammalian brain contains many millions, and in some cases billions, of neurons. No two neurons are identical, yet neurons are regularly categorized into classes and types based on a variety of criteria. This cell type categorization is both logistically and conceptually advantageous for understanding how cells and circuits enable brain function.

Early efforts to define and distinguish groups of neurons focused on the brain areas in which they reside, their precise locations within a given brain area, and/or features of their dendritic, somatic, and/or axonal compartments. Electrophysiological, biochemical, immunohistochemical, and genetic methods have subsequently helped define and distinguish increasingly more specific sets of neurons (reviewed recently in (Klausberger and Somogyi, 2008; Huang and Paul, 2019; Tremblay et al., 2016; Zeng and Sanes, 2017; Cembrowski and Spruston, 2019)). Despite the diversity and amount of data collected and analyzed to this point, the field has not readily converged on a consensus number, or even definition, of neuronal cell types (Klausberger and Somogyi, 2008); indeed, efforts to determine a consensus number of cell types have been limited by well-documented challenges (Ascoli et al., 2008; DeFelipe et al., 2013; Tebaykin et al., 2018).

Recent advances in the collection and analysis of single cell RNA-sequencing (scRNAseq) data provide a promising new way to define cell types and how they relate to one another (Saunders et al., 2018; Tasic et al., 2018; Zeisel et al., 2018; Zeisel et al., 2015; Tasic et al., 2016; Shekhar et al., 2016). This relatively new, transcriptomic approach to neuronal classification is powerful because quantitative expression data from thousands of genes is obtained from each neuron (i.e., high dimensionality) and scRNAseq data can be collected from thousands to millions of individual cells (i.e., high scalability). Moreover, new methods such as the Patch-seq recording technique (Fuzik et al., 2016; Cadwell et al., 2016; Scala et al., 2019; Fö ldy et al., 2016) enable one to directly relate the transcriptomic features of a given neuron to other identifying characteristics of the same neuron - e.g., the neuron’s precise location, morphology, and/or intrinsic electrophysiological properties.

Two of our previous efforts have systematically defined neuronal cell types in mouse visual cortex based on either transcriptomic characteristics using a high-throughput scRNAseq pipeline (Tasic et al., 2018; Tasic et al., 2016) or, from a separate standardized pipeline, electrophysiological and morphological characteristics (Gouwens et al., 2019). In this study we endeavored to classify neuronal types in a more integrated manner by capturing the transcriptomic, electrophysiological and morphological properties of individual neurons via the Patch-seq technique. We chose to focus our initial efforts on cortical GABAergic interneurons for several reasons. First, the morphology of GABAergic interneurons is better preserved in ∼350 um thick *in vitro* slice preparations than that of glutamatergic pyramidal neurons with known long-range projections. Second, transcriptomically-defined types of GABAergic interneurons are more readily conserved across cortical brain areas (Tasic et al., 2018). Third, the number of GABAergic cell types estimated from our scRNASeq data (Tasic et al., 2018) and morpho-electric data (Gouwens et al., 2019) collected in mouse primary visual cortex (VISp)i.e., the number of VISp GABAergic “t-types” (60) and “me-types” (26) differs by more than a factor of two. Such discrepancies are not unique to VISp; studies applying anatomical, physiological, and neurochemical approaches in other cortical areas have also arrived at varying numbers of GABAergic neuronal types (Tremblay et al., 2016; Jiang et al., 2015; Markram et al., 2015). Resolving these discrepancies and deriving a unified set of GABAergic cell types based on congruence among different modalities will provide an essential foundation for achieving a common understanding of the function of these cell types across the field.

Evidence collected to date strongly supports the presence of four major subclasses of cortical GABAergic neurons: parvalbumin (Pvalb)-positive, somatostatin (Sst)-positive, vasoactive intestinal peptide (Vip)positive cells, and cells that express 5-hydroxytryptamine receptor 3A but lack Vip (Htr3a+/Vip-) (Tremblay et al., 2016). Each of these subclasses can be subdivided into several types, largely defined by their axonal morphologies. For example, the Pvalb subclass can be split into basket (and related) cells and chandelier cells; the Sst subclass into Martinotti cells and non-Martinotti cells; the Vip subclass into bipolar and multipolar cells; and the Htr3a+/Vip- subclass into neurogliaform cells and single bouquet cells. Our recent single-cell transcriptomic study of two mouse cortical areas (Tasic et al., 2018), VISp and anterolateral motor cortex (ALM), confirmed these four major subclasses and revealed two additional subclasses, named Sncg and Serpinf1, that are related to the Vip and Htr3a+/Vip(renamed to Lamp5) subclasses.

At the finest (terminal) level of our transcriptomic cell type taxonomy, we identified 60 GABAergic transcriptomic types altogether, each belonging to one of the subclasses. Furthermore, using highly specific transgenic driver lines, we were able to correlate some transcriptomic types (t-types) with some well-known or newly defined morpho-electric types (me-types) (Tasic et al., 2018; Gouwens et al., 2019); for example, the Pvalb Vipr2 and Sst Chodl t-types correspond to chandelier cells and *Nos1*+ long-projecting Sst neurons, respectively. The morpho-electric profiles of most t-types, however, are largely unknown and their correspondence to historically defined GABAergic types has not been established.

Here we report results obtained from the analysis of over 3,700 standardized, quality-controlled Patchseq recordings from GABAergic interneurons in visual cortex of the mature mouse. The dataset, to be made freely available to the public in June of 2020, facilitates answers to a variety of critical questions, including: (1) to what extent do neurons categorized into the same types on the basis of scRNASeq data (i.e., within each t-type) exhibit similar morphological and/or electrophysiological features; (2) to what extent do neurons belonging to different t-types exhibit distinct electrophysiological and/or morphological features; (3) to what extent do transcriptomic and morpho-electric features covary, and are thus mutually predictable; and (4) can a finite number of cell types be defined with maximal congruence across modalities (or are cell “types” infinitely divisible, or too continuous to subdivide)? Answering these and other related questions is a prerequisite to interpreting high-throughput molecular surveys of the brain via FISSEQ (Lee et al., 2014), MERFISH (Chen et al., 2015), SeqFISH (Coskun and Cai, 2016), MAPSeq (Kebschull et al., 2016), osmFISH (Codeluppi et al., 2018) and conceptually similar approaches. The answers will also help define neuronal cell types and features that differ between them, reveal the nature of cellular diversity and its underlying rules, and ultimately improve our understanding of cellular and circuit functions. In addition, the increased inhibitory neuron diversity accessed through transcriptomic types may reveal new insights about structural and electrophysiological organization of mouse visual cortex.

## Results

To establish direct correspondence between the electrical, morphological, and transcriptomic features of mouse cortical neurons, we built a standardized pipeline to extract high quality data from all three modalities from single cells (see Figure 1A, Figure S1, and Methods). In particular, using a modified version of the Patch-seq technique (Cadwell et al., 2016; Fuzik et al., 2016; Fö ldy et al., 2016), we characterized electrophysiological responses to a series of hyperpolarizing and depolarizing current injections, extracted and reverse transcribed nuclear and cytosolic mRNA, sequenced the resulting cDNA using the same scRNAseq method, SMART-Seq v4, as in our transcriptomic study (Tasic et al., 2018), and mapped each cell’s soma location and cortical depth into the Allen Mouse Common Coordinate Framework (CCF) (Kuan et al., 2015). Both the transcriptomic and electrophysiological data from 3,708 cortical GABAergic interneurons passed our QC criteria (see Methods and Figure S1A); the 2341 cells that exhibited adequate biocytin filling were imaged at 63X resolution, and the dendritic and axonal morphologies of 350 of these cells were reconstructed.

**Figure 1:**
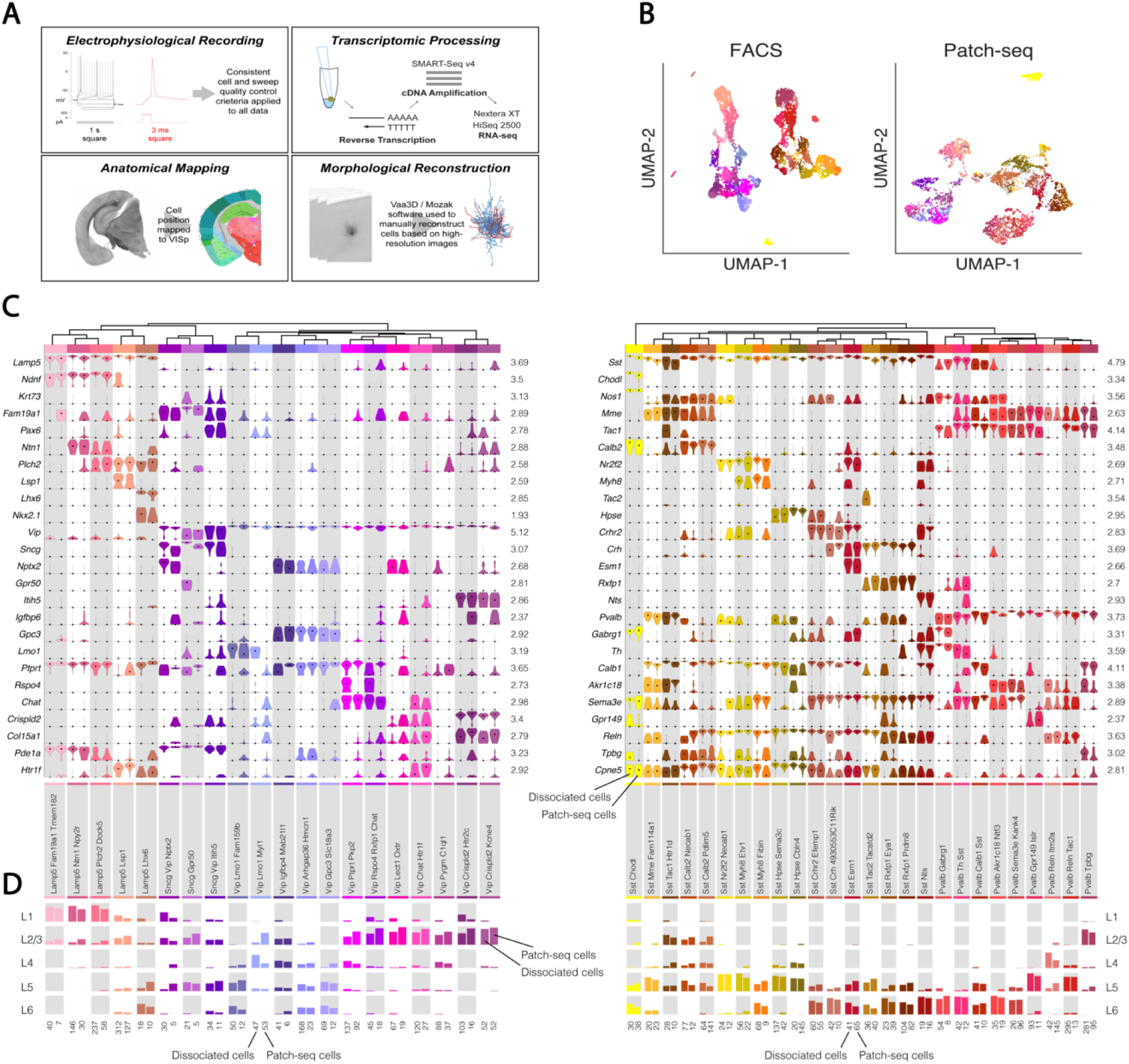
Transcriptomic analysis of scRNASeq data obtained from Patch-seq recordings. (A) Summary of major steps of the Patch-seq data collection and processing pipeline. (B) UMAP plots based on 40 (FACS) or 20 (Patch-seq) principal components derived from log_2_ normalized CPM (counts per million) values of gene expression of the 4,020 most differentially expressed genes across neuron types in the reference data set (n=6,080 dissociated cells, left, and n=3,417 cells from Patch-seq recordings, right). 2,760 “highly consistent” cells (see Figure S1G) are shown in color, 657 additional “moderately consistent” cells are shown in gray. (C) Marker gene expression distributions within each t-type are represented by pairs of violin plots corresponding to Patch-seq recordings and dissociated cells from (Tasic et al., 2018) (right and left in each column for each type, respectively). Rows are genes, black dots are medians. Values within each row are normalized between 0 and maximum detected (shown on the y axis), displayed on a log_10_ scale. (D) Layer distribution of cells for each t-type in both datasets. In each column the dissociated cells and Patch-seq recordings are shown on the left and right, respectively. The layer information for dissociated cells was inferred from layer-enriching dissections. The location of the cells assayed via Patch-seq recordings was determined by pinning neuronal soma location to the common coordinate framework (see Methods). Total number of cells from Patch-seq recordings and dissociated cells in each type are shown below each column on right and left, respectively. For (C-D), only cells from transgenic lines common to both data sets (see Figure S4) and only types with more than 5 cells, were used (n = 4,117 dissociated cells and n=1,790 Patch-seq recordings).

### Relating neurons assayed via Patch-seq recordings to transcriptomic types

The scRNAseq transcriptomes from cDNA obtained from Patch-seq recordings of individual GABAergic interneurons exhibited a similar number of total genes as the scRNAseq transcriptomes obtained from individual dissociated neurons in our previous transcriptomic study (Figure S2A). Moreover, the two transcriptomic datasets exhibited a similar number of discrete clusters when visualized via a Uniform Manifold Approximation and Projection (UMAP, (Becht et al., 2018)) method (Figure 1B). However, we (Figure S2B) and others (Tripathy et al., 2018) found that the mRNA/cDNA collected from one neuron via a Patchseq recording often includes contaminating mRNA/cDNA from adjacent neurons and/or non-neuronal cells. Also, cDNA collected from Patch-seq recordings is more likely to exhibit dropout — missing gene expression due to stochastic failure to capture mRNA from single cells/single nuclei (Figure S2B).

To leverage the quality and large size of transcriptomic datasets obtained from dissociated neurons, we chose to map transcriptomes obtained via Patch-seq recordings to transcriptomic types (t-types) derived from analysis of cDNA obtained from dissociated cells (Tasic et al., 2018). This mapping procedure was designed to minimize the contribution of contamination from non-neuronal cells (such as glia) and nearby excitatory neurons. The steps included first preparing a VISp-specific hierarchically-organized reference tree (based on transcriptomes from dissociated cells from VISp of our previous transcriptomic study), which contained 93 t-types, and choosing marker genes for the mapping i.e., the genes that best distinguished groups at each branch of the tree. Next, we mapped Patch-seq transcriptomes to the reference tree via a bootstrap approach in which only 70% of the available genes and reference cells were used in each of 100 iterations.

We applied additional filters after the initial mapping (see Figure S1 and Methods). One particularly important filter focused on the confidence with which Patch-seq transcriptomes mapped to one or more reference t-types. The transcriptomic cell type taxonomy of (Tasic et al., 2018) found that some t-types were closely related to each other with many cells having as “intermediate” identity between two types. Other types were highly distinctive from each other with no confusion. However, due to contamination and dropout of expressed genes, some Patch-seq transcriptomes unexpectedly presented ambiguous identities even among highly distinct types. To distinguish between “expected” and “unexpected” ambiguity in mapping between t-types, we computed the reference mapping probability matrix between t-types (defined as the probability of a cell type being confused with other cell types in the reference dissociated cell data set). We then used Kullback-Leibler (KL) divergence between the mapping probability distributions and the reference mapping probability distribution as measure of discrepancy. Cells assayed via Patch-seq recordings with divergence greater than 2 were considered “inconsistent” cells. For the remaining cells, the correlation between their gene expression and the average gene expression of all the cell types was computed; cells with correlations below 0.5 were also labeled as “inconsistent.” For the rest of the cells, the mapping quality was determined by bootstrapping confidence. We found that 3,417 of the 3,708 cells (2,760 “highly consistent” and 657 “moderately consistent” cells in Figure S1G) were mapped to one or more than one type with more than 70% confidence (Figure S3, Methods). When we performed analyses by t-type, we restricted the data set to the 2,760 cells with highly-consistent mapping to limit the effect mapping ambiguities due to technical issues on our results.

Several results suggested that this mapping approach is meaningful. First, an independent approach that did not rely on the hierarchical structure of the transcriptomic tree assigned cells to similar t-types (Figure S5). Second, cells obtained from transgenic driver mice that label subclasses or specific types of GABAergic interneurons map to similar t-types regardless of whether the examined transcriptomes originated from dissociated cells or Patch-seq recordings (Figure S4). Third, the relative expression level of key marker genes was similar in cells from the two data sets assigned to the same t-type (Figure 1C). Fourth, the layer distribution of cells from Patch-seq recordings that mapped with high confidence to a given ttype closely resembled the layer distribution of dissociated cells mapping to the same t-type (Figure 1D).

Alignment of our Patch-seq recordings to the CCF enabled detailed insight into the cortical areas and cortical layers from which scRNAseq data were obtained. For example, 83% of our Patch-seq recordings were performed in VISp while the rest were in surrounding higher visual areas (Figure 2A). Moreover, while data obtained from dissociated cells indicated a laminar preference for multiple t-types (see Figure 1D), the additional anatomical precision gained from targeting individual cells with recording electrodes in brain slices revealed further layer specificity (Figure 2B). For example, we found that some Sst t-types in L5 were preferentially located in the upper part (e.g., Sst Calb2 Pdlim5, Sst Hpse Cbln4, Sst Hpse Sema3c), while others occupied the lower part of L5 (e.g., Sst Esm1, Sst Rxfp1 Eya1, Sst Crhr2 Efemp1). Similarly, some t-types were found right at the L1-L2/3 border (Vip Col15a1 Pde1a, Pvalb Vipr2) while others were distributed across L2/3 (Vip Chat Htr1f, Pvalb Tpbg). A smaller number of t-types were notably found across multiple layers (e.g., Lamp5 Lsp1, Sncg Vip Itih5, and Sst Calb2 Pdlim5). The specific laminar (or sublaminar) locations of many GABAergic t-types across subclasses provides a strong anatomical correlate of the cell groupings defined based on transcriptomic data alone.

**Figure 2:**
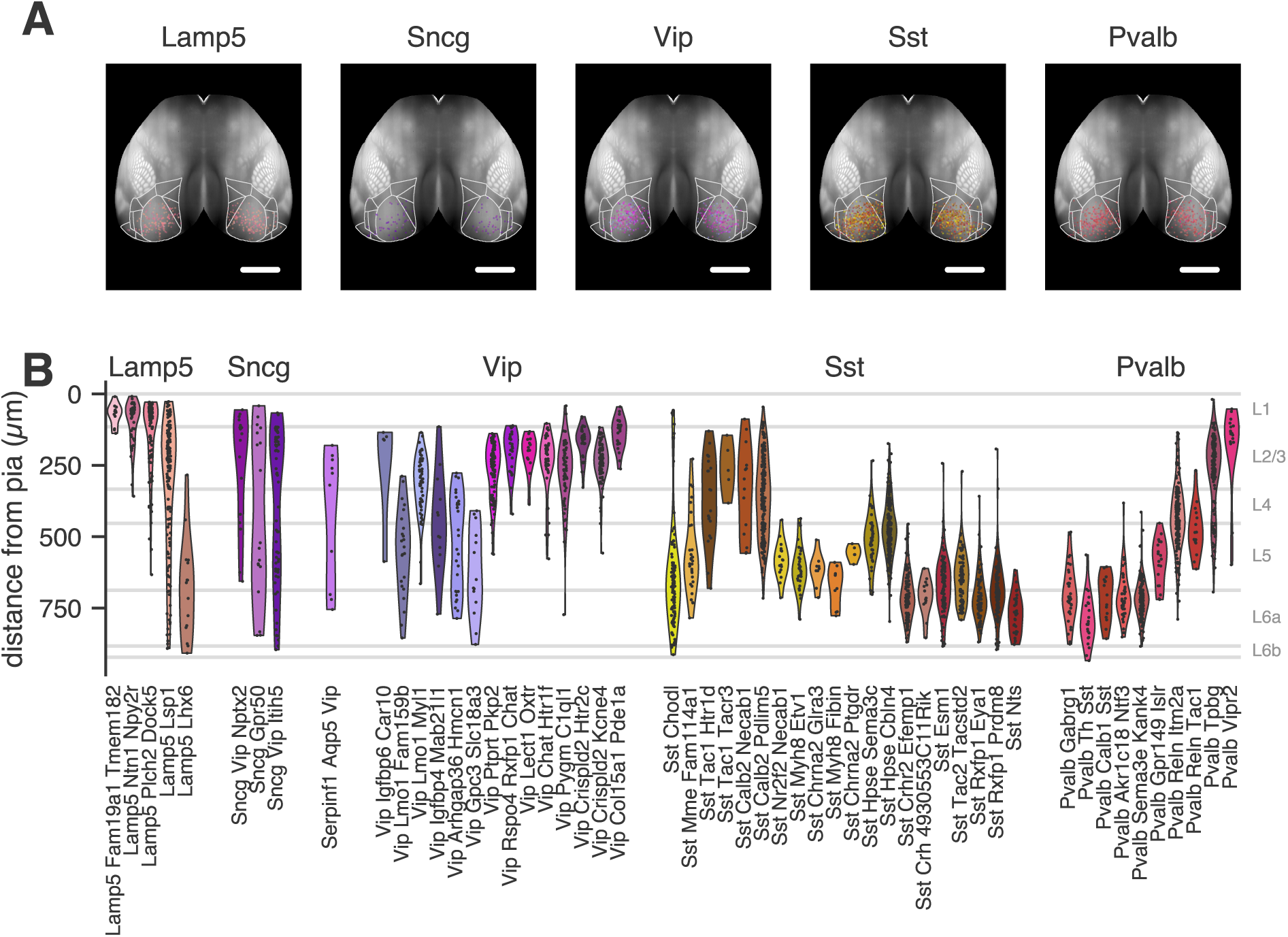
Positions of cells from different t-types in a common reference space. (A) Top-view projections of positions of recorded cells (n=2,739 cells) by transcriptomic subclass and t-type (colors). Lines indicate borders of visual cortical areas. Scale bar: 2 mm. (B) Depth from pia by t-type (n=2,739 cells) where positions are aligned to average cortical layer thicknesses. Only t-types with at least 5 highly-consistent mapped cells are shown in (A) and (B). were spread across spaces occupied by different transcriptomic subclasses. Most t-types appeared to have consistent electrophysiological features, but in principle the features of a given t-type could be essentially the same as another. To test whether this was the case, we fit statistical models of pairs of t-types with either two components (one for each t-type) or a single component after merging the t-types together (Methods). We then assessed how much better the two-component model described the data than the single-component model by calculating the log-likelihood ratio (LLR) between the models (Figure 3E). Most pairs were significantly better fit by separate distributions for each t-type rather than by a single distribution describing the pair, indicating that most t-types systematically differed from each other in their electrophysiological properties. Most of the non-significant differences were found between related pairs of t-types, such as between Sst Myh8 Etv1 and Sst Myh8 Fibin, which are the terminal leaves of the same branch of the transcriptomic tree (see Figure 1C).

### Characterizing the intrinsic electrophysiological features of transcriptomic types

How homogenous are the electrophysiological properties of neurons within a given t-type? How distinguishable are the electrophysiological properties of cells mapped to different t-types? To begin answering these questions, we analyzed all 3,708 neurons’ electrophysiological responses to a standardized currentclamp protocol composed of square-pulse current injections (see Figure S6 for example responses by t-type). We calculated 12 “feature vectors,” capturing properties such as action potential shape and normalized response to hyperpolarizing current indicative of sag potential (Figure 3A), from various aspects of the electrophysiological data. Applying a sparse principal components analysis (sPCA) to each feature vector aggregated the most informative sparse principal components (sPCs, see Methods, (Gouwens et al., 2019)). Projecting the sPCs onto two dimensions using the UMAP method provided a useful visualization of electrophysiological feature gradients across all 3,708 cells (Figure 3B).

**Figure 3:**
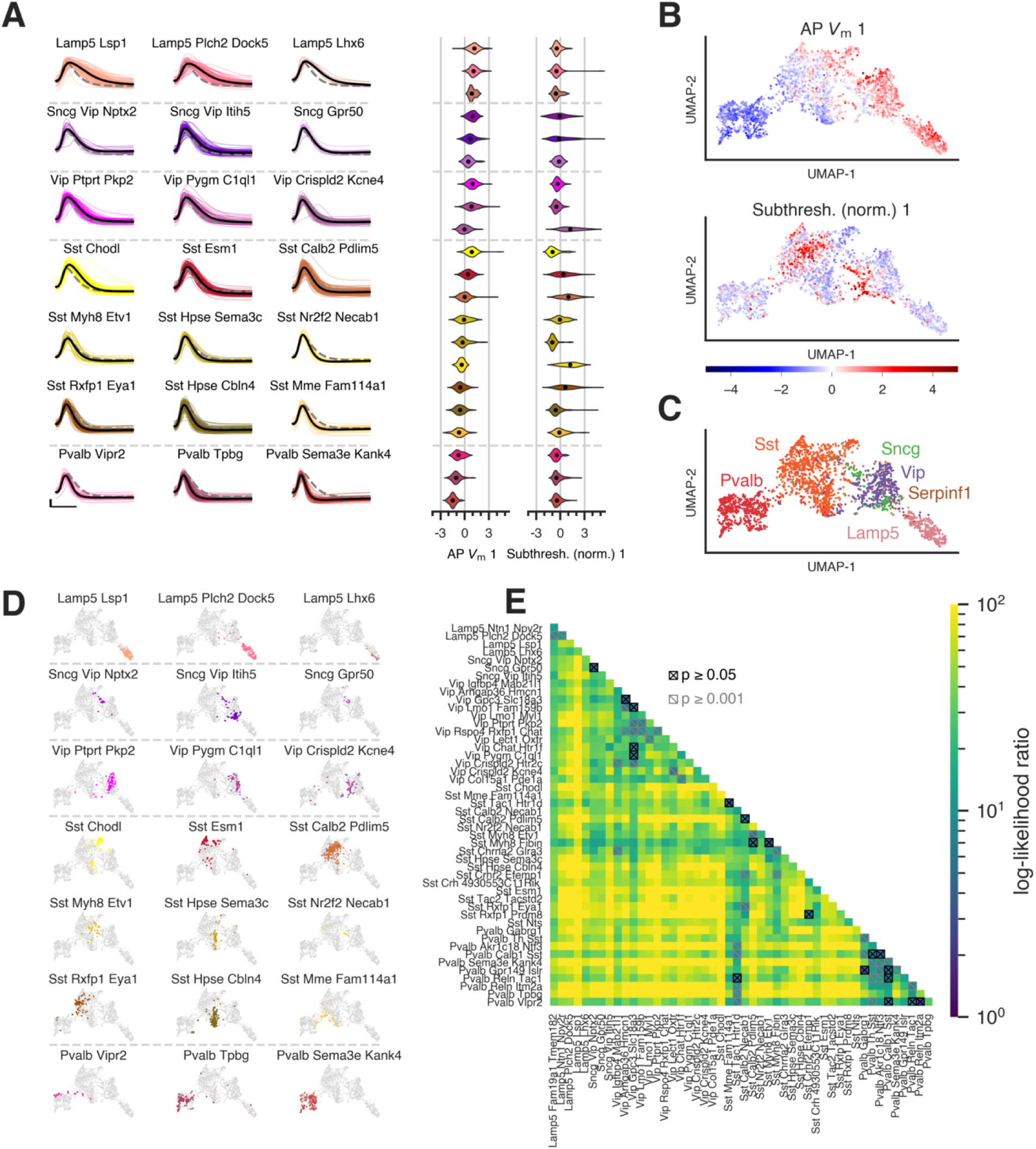
Electrophysiological characterization of transcriptomic types. (A) First action potentials evoked by “long square” (1 s) current injection for cells from 21 example transcriptomic types (ttypes) from Lamp5, Sncg, Vip, Sst, and Pvalb subclasses. Individual cell traces (colors), mean traces for each t-type (black lines) and overall mean (dashed gray line) are shown. Scale bar: vertical 25 mV, horizontal 1 ms. Violin plot of two z-scored sparse principal components (“AP *V*_m_ 1,” “subthresh. norm. 1”) across example t-types. Dashed horizontal lines separate t-types from different subclasses. T-types are sorted by median “AP *V*_m_1” values within each subclass. (B) UMAP plots using 44 z-scored sparse principal component values collected from all 12 electrophysiology feature vectors. “AP *V*_m_ 1” (top) and “subthresh. norm. 1” (bottom) values are shown for each of 3,708 cells (color). (C-D) UMAP plots with 2,760 cells (“highly consistent cells,” see Figure S1G) labeled by transcriptomic subclass (C) and by example t-types (D). Dashed horizontal lines separate t-types from different subclasses. (E) Heatmap of log-likelihood ratios indicating the improvement gained by fitting each of a pair of t-types separately vs together (lighter colors indicate greater improvement). The statistical significance of the improvement was assessed by parametric bootstrap; pairs that do not exceed statistical significance levels of p=0.05 and p=0.001 are indicated by crosses and slashes, respectively. Only t-types with at least 10 highly-consistent mapped cells are shown in (E).

Examining the location of the 2,760 cells with “highly consistent” t-type mapping demonstrated that, at the subclass level, most GABAergic interneurons occupied one of 4 large clouds in the electrophysiology UMAP projection, corresponding to the Lamp5, Vip, Sst and Pvalb transcriptomic subclasses (Figure 3C). Cells of the Sncg and Serpinf1 subclasses fell into smaller regions near the Vip cells. Cells mapping to individual t-types were typically found in a similar location (Figure 3D, Figure S7), suggesting some degree of consistency in their electrophysiological properties. However, this consistency varied across t-types, with some having very tight distributions (e.g., Sst Nr2f2 Necab1), while others (e.g.,Sst Mme Fam114a1)

### Characterizing the morphological features of transcriptomic types

For a subset (350 cells) of transcriptomically and electrophysiologically characterized neurons from the Sst, Pvalb, Vip, Sncg, and Lamp5 subclasses, we also performed morphological analysis. Example mor-phologies from selected t-types are shown aligned to an average cortical template (see Methods), followed by soma and averaged dendrite and axon distributions with respect to cortical layers (Figure 4A; all reconstructions can be found in Figures S8-S11). Example neurons were chosen by calculating a pairwise similarity score for axonal branch structure and location (NBLAST score) (Costa et al., 2016; Kohl et al., 2013) for all reconstructed neurons in a t-type. The five neurons with the highest NBLAST scores are shown left to right in descending order, with the leftmost morphology being most characteristic.

**Figure 4:**
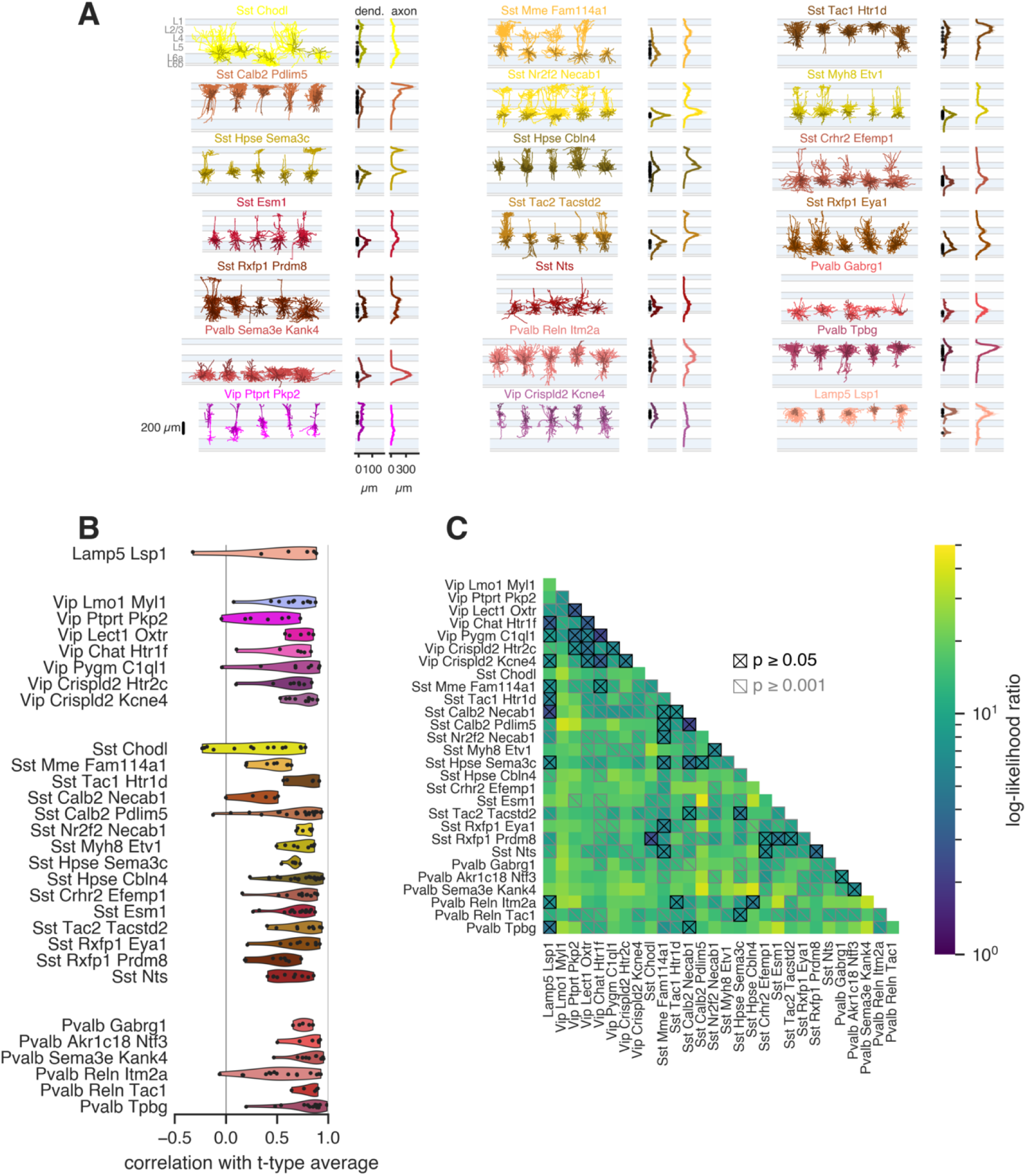
Morphological characterization of transcriptomic types. (A) Representative morphological reconstructions from t-types (selected by NBLAST similarity scores, Methods). Dendrite and axon depth histograms calculated from all reconstructions of the t-type are shown to the right, along with soma depth positions (black dots). Dendrites are in darker colors, axon in lighter colors. Histograms are shown as mean (lines) ± SEM (shaded regions). (B) Correlations between individual cell axon depth histograms and the average histogram of its t-type (excluding itself). (C) Heatmap of log-likelihood ratios indicating the improvement gained by fitting each of a pair of t-types separately vs together (lighter colors indicate greater improvement). The statistical significance of the improvement was assessed by parametric bootstrap; pairs that do not exceed statistical significance levels of p=0.05 and p=0.001 are indicated by crosses and slashes, respectively. Only t-types with at least 5 highly-consistent mapped cells are shown in (B) and (C).

Across subclasses, we observed t-types with distinct dendrite and axon distribution patterns. Specifically, many t-types were distinguished by a layer-selective axonal projection. For example, in the Sst subclass, the Sst Calb2 Pdlim5 type had a L1-dominant (Martinotti-type) axon (DeFelipe et al., 2013), while Sst Tac1 Htr1d had a L2/3-dominant axon, Sst Hpse Cbln4 had a L4-dominant axon, and Sst Rxfp1 Prdm8 had a dominant L5/L6 axon. Similarly, the Pvalb subclass had t-types that predominantly targeted L2/3 (Pvalb Tpbg), L2/3 and L4 (Pvalb Reln Itm2a), and L5 and L6 (Pvalb Sema3e Kank4, Pvalb Gabrg1). Several other Sst t-types also send axon to L1 (Martinotti-like), but either have axon evenly split across two layers (Mme Fam114a1, Hpse Sema3c) or a dominant L5 axon profile (Myh8 Etv1 and Tac2 Tacstd2). In contrast, t-types in the Vip subclass tended to have dendrites and axons that were relatively evenly distributed across multiple layers. The Lamp5 Lsp1 cells had a more variable axon pattern. Interestingly, layer-selective projection patterns often corresponded to the soma locations of each t-type, though not in all cases. Sst Calb2 Pdlim5 cells and Sst Htrd1 exhibited a reliable axon pattern despite having somas spread across L2/3, L4, and L5.

To test how consistent morphologies were within a t-type, we measured the correlation between an individual cell’s axon histogram and the t-type average (Figure 4B). Most cells were positively correlated, indicating consistent axonal innervation patterns. We also examined differences in morphological features among pairs of t-types using the same LLR analysis used with electrophysiological properties (Figure 4C). Though more divisions fell below the significance levels with morphological compared to electrophysiological properties (possibly due to less statistical power due to fewer samples with morphologies), the correspondences of Sst and Pvalb *t-*types with morphology were especially strong; additional data will be required to map other classes with similar precision.

### Variation and overlap of electrophysiological and morphological features among transcriptomic types

While our LLR analyses indicated that systematic differences in other modalities do exist between t-types, the distributions of these features could still exhibit considerable overlap. Therefore, the assignment of an individual cell to a t-type on the basis of electrophysiological and/or morphological features may be difficult if features vary continuously between t-types. To test how well we can assign t-type based on elec-trophysiological and morphological properties, we trained supervised classifiers to predict t-types based on these data modalities.

We trained random forest classifiers on all 44 electrophysiology sPCs to predict either transcriptomic subclass or t-type. The subclass classifier performed at *>*90% accuracy (Figure 5A), indicating that GABAergic transcriptomic subclasses have distinctive electrophysiological features. However, the t-type classifier had a prediction accuracy of 57% (Figure 5B), and prediction error rates varied considerably across t-types. A t-type classifier trained with morphological features only had an accuracy of 41% (Figure 5C), and the pattern of confused types differed from the electrophysiology classifier. The overall lower accuracy could again be in part due to the smaller available number of cells with morphologies. However, classifier accuracy improved to 57% when electrophysiological features were used together with morphological features (Figure 5D).

**Figure 5:**
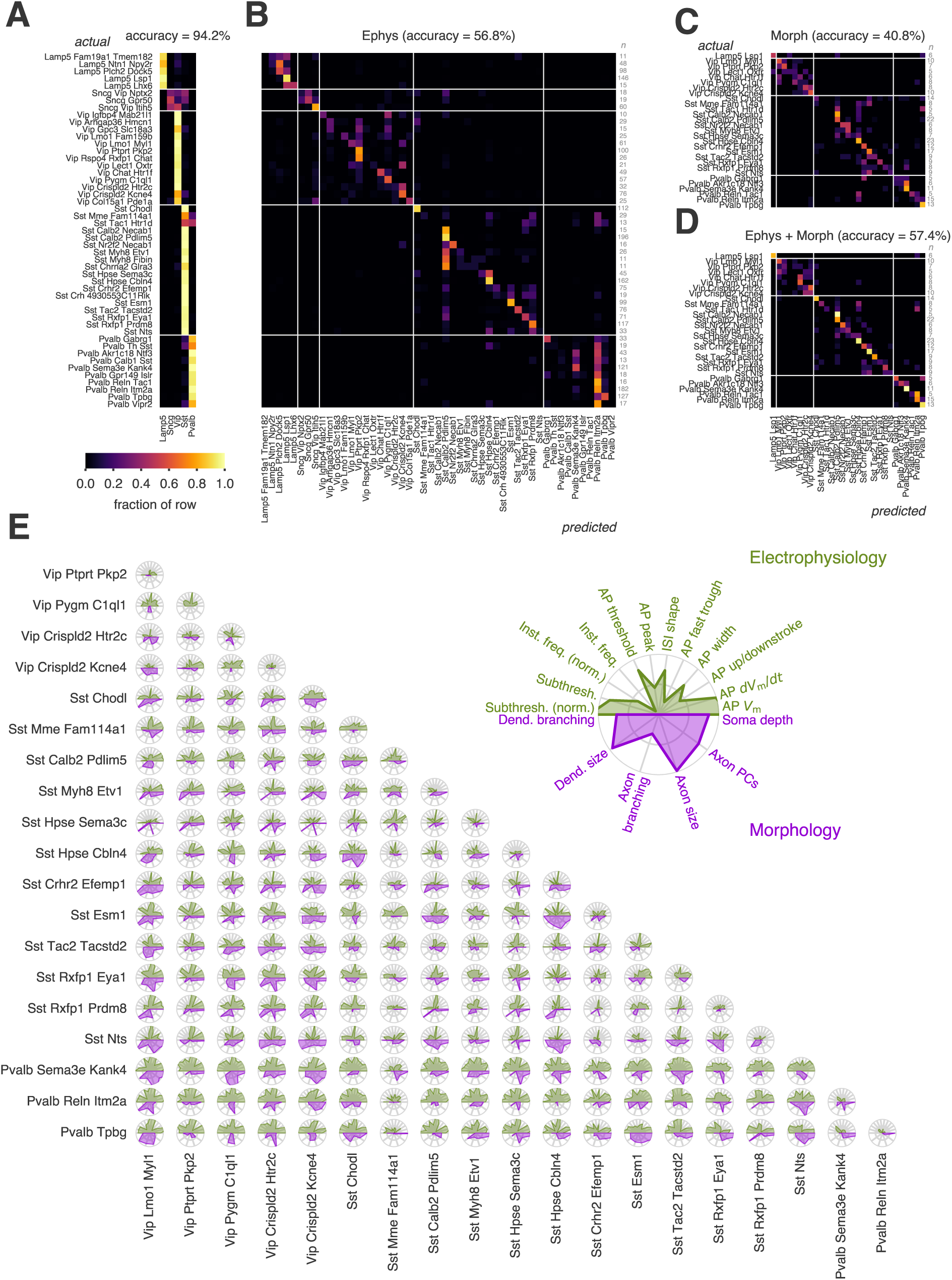
Overlapping properties among different t-types. (A) Out-of-bag confusion matrix of a random forest classifier trained to predict t-subclasses from electrophysiological features. (B) Out-of-bag confusion matrix of a random forest classifier trained to predict t-types from electrophysiological features. (C) Out-of-bag confusion matrix of a random forest classifier trained to predict t-types from morphological features. (D) Out-of-bag confusion matrix of a random forest classifier trained to predict ttypes from combined electrophysiological and morphological features. For (A-D), values are normalized to the row sums (i.e., actual numbers of cells in a given t-type). For (A-B), t-types with at least 10 highly consistent mapped cells each are used. For (C-D), t-types with at least 5 highly consistent mapped cells each are used. (E) Cross-validated *d’* values between pairs of t-types by electrophysiological (green) and morphological (purple) feature groups. Larger *d’* values indicate greater discriminability between pairs of t-types. Only t-types that had at least 20 highly consistent mapped cells with electrophysiology data and seven highly consistent mapped cells with morphological reconstructions are shown. Radial tick interval: 1.

Since classifiers were not able to reliably identify t-types based on electrophysiological and morphological features, and the success rate of predictive classification varied by both t-type and modality, we quantified the degree to which t-types were distinct or overlapping using methods similar to those developed to assess the degree of distinctiveness vs continuity in transcriptomic properties (Harris et al., 2018). For each category of electrophysiological and morphological feature and each pair of t-types, we calculated a cross-validated likelihood ratio statistic (comparing a given held-out cell to either its own class or the other of the pair), then summarized the separation of those likelihood ratios by a *d’* statistic (Figure 5E; higher values indicate more separation).

As expected, closely-related neighboring t-types, such as Sst Rxfp1 Eya1 and Sst Rxfp1 Prdm8, exhibited a high degree of overlap in nearly all feature categories. The Pvalb t-types had a large degree of overlap in electrophysiological features, but had more morphological differences in part due to their different laminar locations (see Figure 2B). Interestingly, the t-type Sst Mme Fam114a1 stood out as being the most similar electrophysiologically to the Pvalb t-types, although it was more separated than the Pvalb t-types were from each other.

### Integrated classification of GABAergic neurons by correspondence between transcriptomic and morpho-electric types

The above analyses revealed that in many cases the cells mapping to a given t-type had consistent electrophysiological and morphological features. However, some t-types had largely overlapping morph-electric features, while others exhibited electrophysiological (e.g., Sst Mme Fam114a1, Sst Tac1 Htr1d) or morphological (e.g., Sst Calb2 Pdlim5, Pvalb Reln Itm2a) heterogeneity. In light of the continuous variation among some t-types, we investigated whether we could group sets of similar t-types that also shared similar electrophysiological and morphological properties. If so, these sets of t-types could be collected into groups that exhibit a high degree of correspondence or congruence across the three modalities assessed here. Therefore, we next performed an unsupervised me-clustering analysis (Gouwens et al., 2019) on the 350 cells with Patch-seq recordings for which we had electrophysiological and morphological data and compared the me-type assignments to the t-types (Figure 6A).

**Figure 6:**
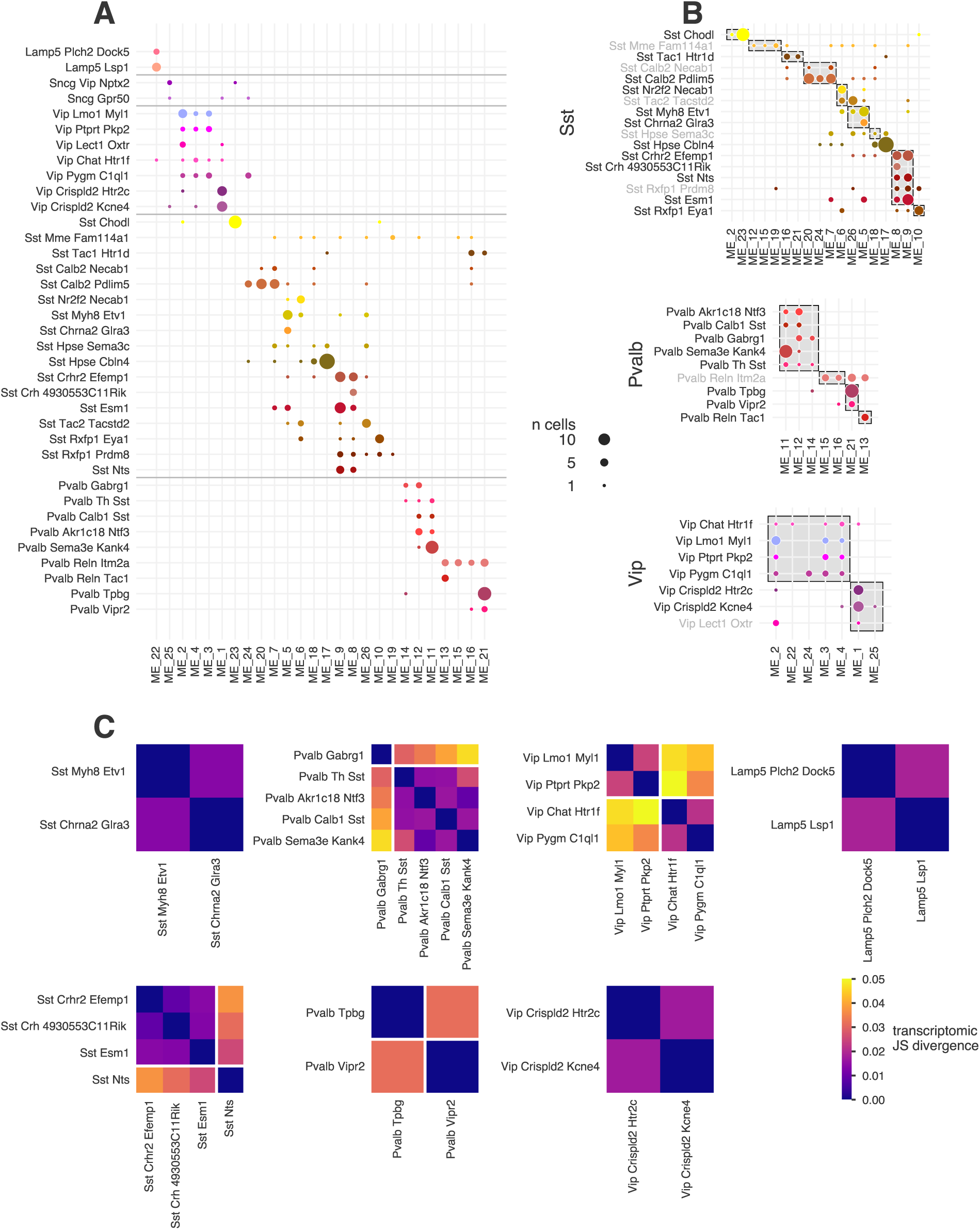
Co-clustering of t-types and me-types. (A) Comparison of t-types with un-supervised joint electrophysiological and morphological clustering results. T-types are ordered by their location on the traxonomy tree. Only t-types with at least 3 highly consistent mapped cells are shown. (B) Spectral co-clustering assignments (gray boxes) by transcriptomic subclass. Gray boxes show the co-cluster assignments. T-types with at least 2/3 of cells within their co-clusters have black text; t-types with less than that have gray text. Size of dot indicates number of cells in (A-B). (C) Transcriptomic similarity (as determined by Jensen-Shannon (JS) divergence) within co-clusters in (B) made up of multiple t-types. Darker colors indicate greater similarity. White lines separate groups with an average between-group JS divergence exceeding 0.03.

Overall, we observed that transcriptomic similarity was closely related to similarity in other modalities, leading to the roughly diagonal correspondence matrix. On the other hand, straightforward one-to-one correspondence was rarely seen. As expected, cells that mapped to different transcriptomic subclasses mapped to mostly non-overlapping sets of me-types. The exceptions included types like Sst Mme Fam114a1 and Sst Tac1 Htr1d which, as mentioned above, showed more Pvalb-like electrophysiological properties. The Sst Tac1 Htr1d t-type had previously been shown to contain cells with gene expression similar to some Pvalb t-types (Tasic et al., 2018). Certain t-types, such as Sst Chodl, Sst Hpse Cbln4, and Pvalb Tbpg, were found mainly in a single respective me-type. For most t-types, however, their cells were distributed across a number of me-types. The reverse was also true; most me-types comprised multiple t-types.

To identify sets of t-types and me-types that corresponded closely to each other, we performed spectral co-clustering (Dhillon, 2001) of t-types and me-types (Figure 6B, gray boxes). Many t-types exhibited good agreement with the co-clusters, defined as having at least two-thirds of their cells within the co-cluster (indicated by black text in Figure 6B), Others, like Sst Mme Fam114a1, had many cells outside those clusters (indicated by gray text in Figure 6B).

For the t-types with good me-type consistency, we asked whether t-types in the same co-cluster were also transcriptomically similar, as determined by their Jensen-Shannon (JS) divergence (Figure 6C; see also Figure S12). Many t-types were quite similar, but some co-clusters had one or more t-types that differed from the others. We used these results to define morphological/electrophysiological/transcriptomic types (met-types) that exhibited a high degree of congruence across all three examined modalities. The co-clusters of Figure 6B were used as a starting point after removing the t-types with poorer agreement (labeled with gray text). Co-clusters were split further if there were large transcriptomic differences among the types. This procedure resulted in twenty identified met-types (Figure 7).

**Figure 7:**
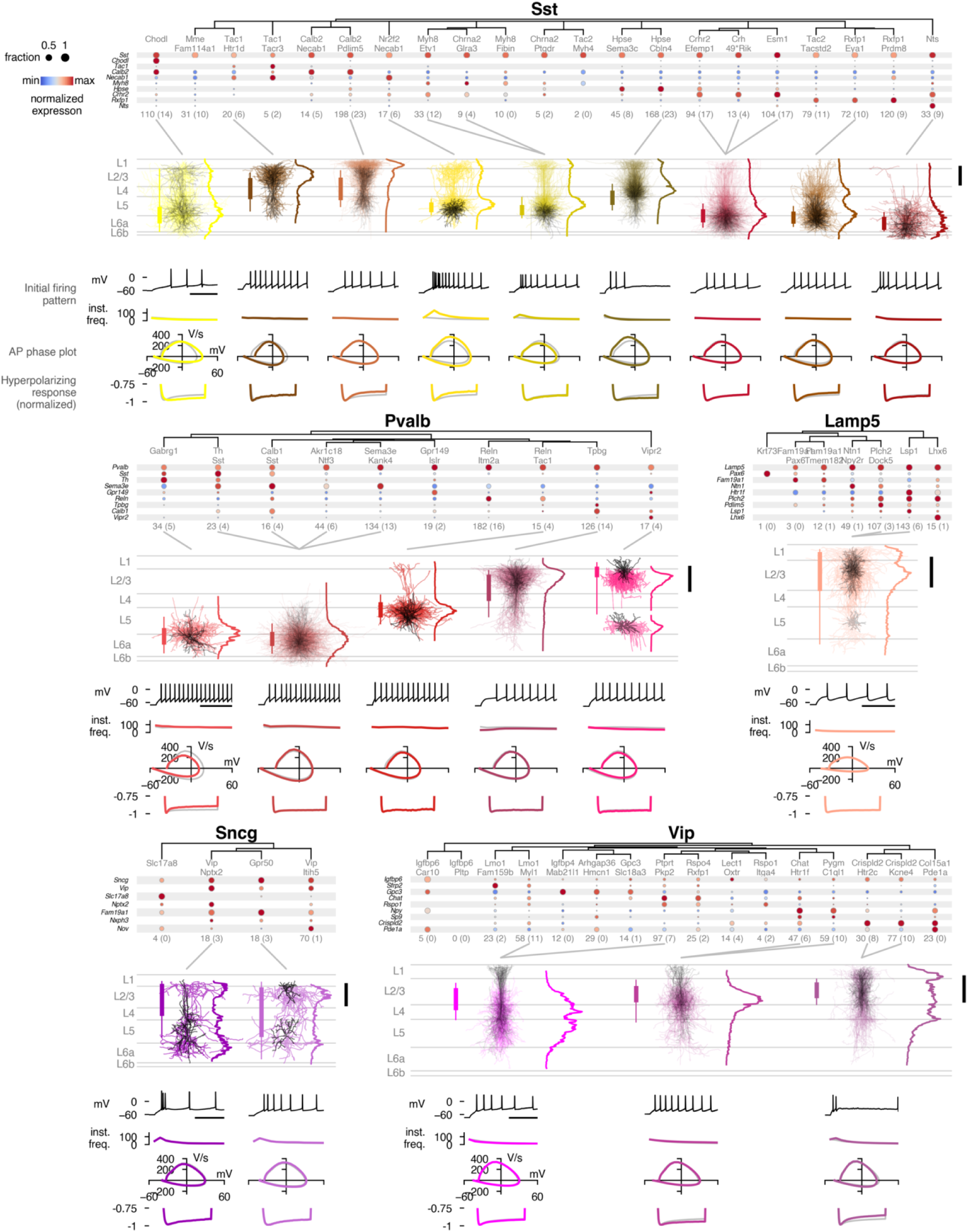
Summary of identified inhibitory met-types. Each transcriptomic subclass is shown with its own dendrogram and with dot plots of example genes that are differentially expressed across the major groups. Expression values (colors) are log-transformed and normalized to the maximum; size of dot indicates fraction of cells expressing the gene within the t-type. Groupings of t-types into “met-types” are shown with gray lines connecting dot plots to morphology plots. Numbers of cells with electrophysiology and a highly-consistent t-type call are below the dot plots. Numbers of cells that also have morphological reconstructions are in parentheses. Morphology plots show all morphologies of the mettypes plotted on top of each other. Soma distributions are indicated by the bars to the left (thicker: 25% to 75% range, thinner: 5% to 95% range). Average axon depth histograms are shown to the right (normalized to the maximum value of each). Scale bar at far right: 200 µm. Electrophysiology summaries include an example trace from a single cell evoked by a rheobase + 40 pA stimulus (scale bar: 100 ms), met-type–wide average instantaneous firing rate (spikes/sec) at that same stimulus amplitude, phase plots of the average first spike of the rheobase trace, and average normalized responses to a hyperpolarizing step. Met-type averages are shown in colors, while subclass-wide averages are shown for comparison in gra

We identified nine met-types within the Sst subclass. The first one (the left-most column in the top row of Figure 7) — mapped to the Sst Chodl t-type, generally occupied deeper layers, and corresponded to non-Martinotti, long-range projecting interneurons (Tasic et al., 2018; Paul et al., 2017; He et al., 2016). The second met-type comprised cells mapping to Sst Tac1 Htr1d with electrophysiological properties most reminiscent of Pvalb interneurons. These cells had APs with low upstroke/downstroke ratios and little axonal innervation of L1. A previous study also observed that cells of this t-type had fast-spiking-like firing properties (Scala et al., 2019). The third met-type contained cells mapping to Sst Calb2 Pdlim5, which were found in cortical layers L2/3 to L5 and exhibited the most exuberant axonal elaborations in L1. As mentioned, this met-type reflects the well-described Martinotti-type neurons, and includes cells that resemble the recently described L2/3 Martinotti and L5/6 fanning-out Martinotti cells (Muñ oz et al., 2017; Nigro et al., 2018). Note that the dominant layer of innervation for the fanning-out type may differ between visual and somatosensory cortex, while overall axon shape remains the same. The fourth met-type contained the cells mapping to Sst Nr2f2 Necab1 that possessed burst-like or very strongly adapting firing patterns; their axons predominantly innervated the region near their somas in L5, with a more minimal projection to L2/3 and superficial L1. This met-type also resembles the L5/6 fanning-out neurons described in these same papers. The fifth met-type (containing cells mapping to Sst Myh8 Etv1 and Sst Chrna2 Glra3) was similar to the previous one but with more variable L1 axon, though its appearance is closer to a T-shaped Martinotti cell (Muñ oz et al., 2017), and its electrophysiological properties are more similar to the Sst subclass average. The sixth met-type mapped to Sst Hpse Cbln4 and was found in L4 and upper L5; it strongly innervated L4 (with a less dominant projection to L1), and exhibited distinct electrophysiological properties including lower AP thresholds, less hyperpolarization-evoked sag, and more transient or irregular firing patterns. This met-type aligns with L5 Non-Martinotti cells in somatosensory cortex (Naka et al., 2019) that have distinct connectivity profiles compared to Martinotti neurons, though in visual cortex these neurons have much more L1 axon. The seventh met-type was composed of cells mapping to the Sst Crhr2 Efemp1, Sst Crh 4930553C11Rik, and Sst Esm1 t-types, was found in deep L5 and L6, and primarily innervated those layers with relatively few axonal projections to L1. The eighth met-type, composed of cells mapping to Sst Rxfp1 Eya1, was similar to the seventh but exhibited more prominent axonal innervation of L4. Neurons with a similar axon innervation pattern (L5/L6 non-Martinottis) have been observed in deep layers of somatosensory cortex (Muñ oz et al., 2017). The cells of the nineth Sst met-type (corresponding to the Sst Nts t-type) were mostly found in L6 and innervated the area around their somas with little projection toward upper layers. The last three Sst met-types had electrophysiological properties closely matching the Sst subclass average.

In this study, we did not attempt to associate t-types with low numbers of samples with an met-type; however, some well-populated Sst t-types were also not associated with an met-type. Sst Mme Fam114a1 was widely spread across a number of me-types (Figure 6B) with little homogeneity in other modalities. Sst Hpse Sema3c contained some cells sharing me-types with Sst Hpse Cbln4 (which it closely resembled transcriptomically), but a number of others were more similar to other Sst L5 types. Likewise, Sst Tac2 Tacstd2 and Sst Rxfp1 Prdm8 had some cells resembling the transcriptomically-similar Sst Rxfp1 Eya1, but the former had many cells with me-types shared with more superficial L5 Sst types, while the latter shared more with deeper L5/L6 types (Figure 6B). Many of these relationships also reflect connections between t-types due to cells with intermediate transcriptomic identities observed by (Tasic et al., 2018).

Five met-types were identified in the Pvalb subclass and mostly reflected different laminar positions. AP shapes were quite consistent across the Pvalb met-types and were typical of a fast-spiking phenotype. The first two met-types were found predominantly in L5 and L6 and had basket-like morphologies with axon and dendrites restricted to L5 and L6. The first contained cells mapped to the transcriptomically-distinct Pvalb Gabrg1 t-type and also exhibited more hyperpolarization-induced sag than the other Pvalb met-types; the second was composed of four Pvalb t-types and contained fast-spiking cells resembling locally-projecting L6 cells (Bortone et al., 2014)and deep basket cells (Markram et al., 2015; Jiang et al., 2015). The third met-type mapped to Pvalb Reln Tac1 and contained mostly L5 fast-spiking cells with dominant L5 axon innervation and a few collaterals extending into superficial L1. The fourth met-type mapped to Pvalb Tpbg and contained mainly L2/3 fast-spiking cells with a dominant L2/3 axonal projection and sparse projections to deeper layers; delays in the onset of AP generation were more common for cells in this met-type. The last met-type (containing Pvalb Vipr2 cells) represented chandelier cells as previously identified (Tasic et al., 2018; Gouwens et al., 2019) and had somewhat more firing frequency adaptation than the other Pvalb groups. Curiously, the Pvalb Reln Itm2a t-type contained the largest number of cells, but its cells exhibited diverse basket-like axonal morphologies, often with translaminar axon branches (Bortone et al., 2014), with somas spanning from L2/3 to L5 and hence corresponding to multiple me-types, including those shared with Pvalb Reln Tac1 and Pvalb Tpbg (Figure 6B). Thus, we did not assign this t-type to a met-type, though we noted it contained most of the L4 fast-spiking basket cells in the data set. This observation, combined with the substantial overlap of electrophysiological properties among the second through fourth Pvalb met-types, suggests a continuous variation of basket and related cells across cortical depth.

Only one met-type could be robustly identified among the Lamp5 cells largely due to the relative dearth of morphological reconstructions. This met-type had APs with high upstroke/downstroke ratios and a neurogliaform-like morphology; the cells were found primarily in L1 and L2/3 but also appeared in deeper cortical layers. Two met-types corresponding to two t-types in the Sncg subclass were identified, one with translaminar dendrites (Sncg Vip Nptx2) and the other with more local, multipolar dendrites (Sncg Gpr50); both were found in most cortical layers. Based on gene expression, Sncg neurons have been linked to *Vip*+/*Cck*+ basket cells (Tasic et al., 2018). We identified three Vip met-types, each including two t-types, all resembling bipolar or bitufted cells (Prö nneke et al., 2015). The first met-type (Vip Lmo1 Myl1 and Vip Ptprt Pkp2) had cells primarily in L2/3 and L4, with dendrites extending into L1, and axon innervating L2/3 through L6. The second met-type (Vip Chat Htr1f and Vip Pygm C1ql1) had a similar morphological profile but with a more concentrated innervation of lower L2/3. The last Vip met-type (Vip Crispld2 Htr2c and Vip Crispld2 Kcne4) innervated all layer, including L1, and had a deeper hyperpolarization-induced sag than the others.

## Discussion

Groups, classes, or types of neurons have often been distinguished on the basis of structural, physiological, biochemical, or genetic data. The (sometimes dramatically) different number of distinguishable neuronal cell types inferred from one data modality versus another, and the challenge of relating groups of neurons distinguished on the basis of one set of cellular attributes to groups of neurons distinguished on the basis of another highlights the power and utility of a more integrated experimental approach. Here, the highthroughput, standardized application of the Patch-seq technique enabled the intrinsic electrophysiological, morphological, and transcriptomic characteristics of individual neurons to be assessed simultaneously.

Electrophysiological and transcriptomic data collected from over 3,700 Patch-seq recordings passed our QC criteria and were mapped to the Allen Mouse Common Coordinate Framework (CCF); scRNASeq data obtained from over 3,400 of these recordings mapped with *>*70% confidence to one or more of the GABAergic transcriptomic types defined in (Tasic et al., 2018). This unprecedented dataset, freely available to the public in June of 2020, enables a broad range of analyses: a major goal of generating, curating, and releasing this resource is to facilitate others’ analyses and interpretations. We hope that detailed analyses of these data will enable individuals familiar with collecting and interpreting one type of data (e.g., electrophysiological data) to better interpret it in the context of other studies.

In the meantime, one major insight gained from our initial analysis of the dataset is that the somas of many GABAergic t-types are restricted to a single cortical layer, one part of that layer, or small regions on either side of a layer boundary (Figure 2) and, in the case of Sst interneurons, elaborate their axons in a dominant cortical layer (Figure 4). Analysis of this dataset supports the presence of nine Sst met-types (Figure 7) — nearly twice as many as described in previous morphological and/or electrophysiological studies (Kawaguchi and Kubota, 1996; McGarry et al., 2010; Markram et al., 2015; Jiang et al., 2015; Muñ oz et al., 2017) — and directly links transcriptomically- and morphologically-defined types of GABAergic neurons. Our data, for example, link a major Sst t-type (Sst Calb2 Pdlim5) with the well-known, frequently-encountered class of neurons referred to as Martinotti cells (Wang et al., 2004; Tremblay et al., 2016; Muñ oz et al., 2017; Scala et al., 2019); neurons mapping to the Sst Calb2 Pdlim5 t-type, are found in L2/3 through L5 and, like classically-defined Martinotti cells, exhibit extensive axon elaborations in L1 relative to other layers. The axons of neurons mapping to all other Sst met-types, by comparison, preferentially arborize outside of L1. Note that all these Sst met-types contain at least one cell that has some degree of L1 projection, and therefore could be considered a Martinotti cell as well; we are not aware of an absolute definition of a Martinotti cell in terms of total amount of axon in L1, though it is generally described as “significant” (Tremblay et al., 2016; Zhou et al., 2020). However, we find that these met-types consistently target one or two other layers to a greater extent than L1. For example, the Sst met-type enriched for faster spiking neurons (Sst Tac1 Htr1d), also located in L2/3 through L5, exhibits dense axonal innervation of L2/3 and largely avoids L1. Sst ttypes found mainly in the deeper part of L5 and in L6 also have dominant L5 and L6 axonal projections and either avoid L4 (Sst Nts) or have axonal enrichment in L4 (Sst Rxfp Eya1). The axons of neurons mapping to the Sst Hpse Cbln4 t-type — one of two major Sst t-types found in L4 — innvervate L1 and, to a greater degree, L4. A dominant L4 projection appears to be a common feature of Sst Hpse Cbln4 neurons across cortical areas (Naka et al., 2019). These results reveal that Sst neurons highly respect laminar boundaries across cortex, they have clear laminar innervation preferences that differ based on their met-type and/or laminar position, and that their dominant laminar target may be a more reliable feature for distinguishing among Sst neurons than the presence or absence of axon in layer 1.

A second major finding is that the electrophysiological (Figure 3) and morphological (Figure 4) properties of neurons mapping to a given t-type are similar to that of other neurons mapping to the same t-type. At the same time, it is frequently difficult to predict the specific t-type of a neuron based on morphological and/or physiological features (Figures 5 and 6). The much lower dimensionality of electrophysiological and morphological data compared to transcriptomic data, and the comparatively small number of morphological reconstructions, may contribute to this disconnect.

Technical aspects of Patch-seq recording, including contamination and dropout, can introduce ambiguity in t-type mapping, so we chose to focus on cells with highly consistent mapping to limit the effects of those issues on our results. However, it is worth noting that some of the ambiguity in types could be biological rather than technical, such that focusing our analysis this way may overly-discretize a landscape with more continuous variation between cells and cell types. Along similar lines, Scala et al. (2020) describe a continuum of variability in the morphology and electrophysiology of glutamatergic and GABAergic neurons in primary motor cortex of the mouse similar transcriptomic cell types frequently exhibit similar morphoelectric features, often without clear boundaries between them. Also, (Que et al., 2020) report that neurons mapping to distinct Pvalb t-types in the mouse hippocampus exhibit similar morphological characteristics.

Altogether, our results support the presence of at least 20 GABAergic interneuron met-types that have congruent transcriptomic, electrophysiological and morphological properties (Figure 7). These met-types represent a version of a unified definition of mouse cortical GABAergic interneuron types backed by multimodal experimental data and enabling cross-modality feature prediction. The number of distinguishable met-types could change in the future for multiple reasons. One is that we do not yet have a sufficiently large number of morphology reconstructions for the Vip, Lamp5 and Sncg subclasses. As a result, it is as yet unknown whether cells mapping to other t-types will resemble the cells in the already-identified met-types, or if they will contribute to new met-types. Moreover, we cannot rule out the possibility that artifacts of the brain slice preparation contribute to or explain observed morphological differences, although the fact that neurons mapping to different met-types (within subclass) exhibit a similar range of total axon lengths suggests that this might not be a major concern.

We also noted that a few GABAergic t-types (e.g., Sst Chrna2 Glra3) that were well-represented by scRNASeq data obtained from dissociated cells (Tasic et al., 2018) appeared relatively infrequently in our data set from Patch-seq recordings; this difference persisted despite using the same, relatively specific transgenic driver lines to label cells under both experimental conditions. Moreover, in several cases, scRNASeq data from PatchSeq recordings mapped more frequently to one t-type (Sst Hpse Cbln4, Pvalb Reln Itm2a) while scRNASeq data from dissociated cells mapped more frequently to another, closely-related t-type (Sst Hpse Sema3c, Pvalb Reln Tac1, respectively). Such discrepancies may reflect gene expression differences in slice versus dissociated cell preparations.

The number of archetypal interneuron groups might also change as additional information about cellular and circuit properties is collected. This could include further refinement of the transcriptomic taxonomy profiled under different states, more detailed electrophysiological and morphological characterization, the addition of proteomic information, local and long-range synaptic connectivity, and (*in vivo*) functional characterization. With the availability of large-scale electromicrograph images, we will soon be able to map morphological phenotypes across datasets to understand the relationship between transcriptomic, morphological and electrophysiological phenotypes and connectivity profiles. In addition, multiplex FISH is already being used to measure the proportions of transcriptomic types within a given brain tissue (Hodge et al., 2019; Moffitt et al., 2018). Combined, these methods will facilitate the development of cell typespecific circuit maps with unprecedented resolution and detail.

## Methods

### Mouse breeding and husbandry

All procedures were carried out in accordance with the Institutional Animal Care and Use Committee at the Allen Institute for Brain Science. Animals (*<*5 mice per cage) were provided food and water ad libitum and were maintained on a regular 12-h light–dark cycle. Animals were maintained on the C57BL/6J background, and newly received or generated transgenic lines were backcrossed to C57BL/6J. Experimental animals were heterozygous for the recombinase transgenes and the reporter transgenes. Standard tamoxifen treatment for CreER lines included a single dose of tamoxifen (40 µL of 50 mg mL^-1^) dissolved in corn oil and administered via oral gavage at P10–P14. Tamoxifen treatment for *Nkx2.1-*CreERT2*;Ai14* was performed at embryonic day (E)17 (oral gavage of the dam at 1 mg per 10 g of body weight), pups were deliv-ered by cesarean section at E19 and then fostered. *Ndnf-*IRES2*-*dgCre animals did not receive trimethoprim induction, since the baseline dgCre activity (without trimethoprim) was sufficient to label the cells with the Ai14 reporter.

### Tissue processing

Mice (male and female) between the ages of P45 and P70 were anesthetized with 5% isoflurane and intracardially perfused with 25 or 50 mL of ice-cold slicing artificial cerebral spinal fluid (ACSF; 0.5 mM calcium chloride (dehydrate), 25 mM D-glucose, 20 mM HEPES buffer, 10 mM magnesium sulfate, 1.25 mM sodium phosphate monobasic monohydrate, 3 mM myo-inositol, 12 mM *N*-acetyl-L-cysteine, 96 mM *N*-methyl-Dglucamine chloride (NMDG-Cl), 2.5 mM potassium chloride, 25 mM sodium bicarbonate, 5 mM sodium L-ascorbate, 3 mM sodium pyruvate, 0.01 mM taurine, and 2 mM thiourea (pH 7.3), continuously bubbled with 95% O_2_/5% CO_2_). Slices (350 µm) were generated (Compresstome VF-300 vibrating microtome, Precisionary Instruments or VT1200S Vibratome, Leica Biosystems), with a block-face image acquired (Mako G125B PoE camera with custom integrated software) before each section to aid in registration to the common mouse reference atlas. Brains were mounted for slicing either coronally or 17° off-coronal to preserve intactness of a neuronal processes and primary visual cortex.

Slices were transferred to an oxygenated and warmed (34 °) slicing ACSF for 10 min, then transferred to room temperature holding ACSF (2 mM calcium chloride (dehydrate), 25 mM D-glucose, 20 mM HEPES buffer, 2 mM magnesium sulfate, 1.25 mM sodium phosphate monobasic monohydrate, 3 mM myo-inositol, 12.3 mM *N*-acetyl-L-cysteine, 84 mM sodium chloride, 2.5 mM potassium chloride, 25 mM sodium bicarbonate, 5 mM sodium L-ascorbate, 3 mM sodium pyruvate, 0.01 mM taurine, and 2 mM thiourea (pH 7.3), continuously bubbled with 95% O_2_/5% CO_2_) for the remainder of the day until transferred for patch-clamp recordings.

### Patch-clamp recording

Slices were bathed in warm (34 °) recording ACSF (2 mM calcium chloride (dehydrate), 12.5 mM Dglucose, 1 mM magnesium sulfate, 1.25 mM sodium phosphate monobasic monohydrate, 2.5 mM potassium chloride, 26 mM sodium bicarbonate, and 126 mM sodium chloride (pH 7.3), continuously bubbled with 95% O_2_/5% CO_2_). The bath solution contained blockers of fast glutamatergic (1 mM kynurenic acid) and GABAergic synaptic transmission (0.1 mM picrotoxin). Thick-walled borosilicate glass (Warner Instruments, G150F-3) electrodes were manufactured (Narishige PC-10) with a resistance of 4–5 Mn. Before recording, the electrodes were filled with ∼1.0-1.5 µL of internal solution with biocytin (110 mM potassium gluconate, 10.0 mM HEPES, 0.2 mM ethylene glycol-bis (2-aminoethylether)-*N*,*N*,*N*’,*N*’-tetraacetic acid, 4 mM potassium chloride, 0.3 mM guanosine 5’-triphosphate sodium salt hydrate, 10 mM phosphocreatine disodium salt hydrate, 1 mM adenosine 5’-triphosphate magnesium salt, 20 µg/mL glycogen, 0.5U/µL RNAse inhibitor (Takara, 2313A) and 0.5% biocytin (Sigma B4261), pH 7.3). The pipette was mounted on a Multiclamp 700B amplifier headstage (Molecular Devices) fixed to a micromanipulator (PatchStar, Scientifica).

The composition of bath and internal solution as well as preparation methods were made to maximize the tissue quality of slices from adult mice, to align with solution compositions typically used in the field (to maximize the chance of comparison to previous studies), modified to reduce RNAse activity and ensure maximal gain of mRNA content.

Electrophysiology signals were recorded using an ITC-18 Data Acquisition Interface (HEKA). Commands were generated, signals processed, and amplifier metadata were acquired using MIES (https://github.com/AllenInstitute/MIES/), written in Igor Pro (Wavemetrics). Data were filtered (Bessel) at 10 kHz and digitized at 50 kHz. Data were reported uncorrected for the measured (Neher, 1992) –14 mV liquid junction potential between the electrode and bath solutions.

Prior to data collection, all surfaces, equipment and materials were thoroughly cleaned in the following manner: a wipe down with DNA away (Thermo Scientific), RNAse Zap (Sigma-Aldrich) and finally with nuclease-free water.

After formation of a stable seal and break-in, the resting membrane potential of the neuron was recorded (typically within the first minute). A bias current was injected, either manually or automatically using algorithms within the MIES data acquisition package, for the remainder of the experiment to maintain that initial resting membrane potential. Bias currents remained stable for a minimum of 1 s before each stimulus current injection.

To be included in analysis, a cell needed to have a *>*1 Gn seal recorded before break-in and an initial access resistance *<*20 Mn and *<*15% of the Rinput. To stay below this access resistance cut-off, cells with a low input resistance were successfully targeted with larger electrodes. For an individual sweep to be included, the following criteria were applied: (1) the bridge balance was *<*20 Mn and *<*15% of the *R*_input_; (2) bias (leak) current 0 ± 100 pA; and (3) root mean square noise measurements in a short window (1.5 ms, to gauge high frequency noise) and longer window (500 ms, to measure patch instability) *<*0.07 mV and 0.5 mV, respectively.

mUpon Extracting the nucleus at the conclusion of the electrophysiology experiment led to a substantial increase in transcriptomic data quality and abilitUpon completion of electrophysiological examination, the pipette was centered on the soma or placed near the nucleus (if visible). A small amount of negative pressure was applied (∼-30 mbar) to begin cytosol extraction and attract the nucleus to the tip of pipette. After approximately one minute, the soma had visibly shrunk and/or the nucleus was near the tip of the pipette. While maintaining the negative pressure, the pipette was slowly retracted in the x and z direction. Slow, continuous movement was maintained while monitoring pipette seal. Once the pipette seal reached *>*1Gn and the nucleus was visible on the tip of the pipette, the speed was increased to remove the pipette from the slice. The pipette containing internal solution, cytosol and nucleus was removed from pipette holder and contents were expelled into a PCR tube containing the lysis buffer (Takara, 634894).

### cDNA amplification and library construction

We used the SMART-Seq v4 Ultra Low Input RNA Kit for Sequencing (Takara, 634894) to reverse transcribe poly(A) RNA and amplify full-length cDNA according to the manufacturer’s instructions. We performed reverse transcription and cDNA amplification for 21 PCR cycles in 8-well strips, in sets of 12–24 strips at a time. At least 1 control strip was used per amplification set, which contained 4 wells without cells and 4 wells with 10 pg control RNA. Control RNA was either Mouse Whole Brain Total RNA (Zyagen, MR-201) or control RNA provided in the SMARTSeq v4 kit. All samples proceeded through Nextera XT DNA Library Preparation (Illumina FC-131-1096) using Nextera XT Index Kit V2 Set A (FC-131-2001). Nextera XT DNA Library prep was performed according to manufacturer’s instructions except that the volumes of all reagents including cDNA input were decreased to 0.4× or 0.5× by volume. Subsampling was conducted to a read depth of 0.5 million per cell.

### Sequencing data processing

Fifty-base-pair paired-end reads were aligned to GRCm38 (mm10) using a RefSeq annotation gff file retrieved from NCBI on 18 January 2016 (https://www.ncbi.nlm.nih.gov/genome/annotationeuk/all/). Sequence alignment was performed using STAR v2.5.3 (Dobin et al., 2013) in two pass Mode. PCR duplicates were masked and removed using STAR option “bamRemoveDuplicates.” Only uniquely aligned reads were used for gene quantification. Gene counts were computed using the R Genomic Alignments package summarize (Lawrence et al., 2013) Overlaps function using “IntersectionNotEmpty” mode for exonic and intronic regions separately.

### Morphological reconstruction

#### Biocytin histology

A horseradish peroxidase (HRP) enzyme reaction using diaminobenzidine (DAB) as the chromogen was used to visualize the filled cells after electrophysiological recording, and 4,6-diamidino-2-phenylindole (DAPI) stain was used identify cortical layers as described previously (Gouwens et al., 2019).

#### Imaging

Mounted sections were imaged as described previously (Gouwens et al., 2019). Briefly, operators captured images on an upright AxioImager Z2 microscope (Zeiss, Germany) equipped with an Axiocam 506 monochrome camera and 0.63x optivar. Two-dimensional tiled overview images were captured with a 20X objective lens (Zeiss Plan-NEOFLUAR 20X/0.5) in brightfield transmission and fluorescence channels. Tiled image stacks of individual cells were acquired at higher resolution in the transmission channel only for the purpose of automated and manual reconstruction. Light was transmitted using an oil-immersion condenser (1.4 NA). High-resolution stacks were captured with a 63X objective lens (Zeiss Plan-Apochromat 63x/1.4 Oil or Zeiss LD LCI Plan-Apochromat 63x/1.2 Imm Corr) at an interval of 0.28 µm (1.4 NA objective) or 0.44 µm (1.2 NA objective) along the Z axis. Tiled images were stitched in ZEN software and exported as single-plane TIFF files.

#### Anatomical location

To characterize the position of biocytin-labeled cells in the mouse brain, a 20x brightfield and/or fluorescent image of DAPI (4’,6-diamidino-2-phenylindole) stained tissue was captured and analyzed to determine layer position and region. Soma position of reconstructed neurons was annotated and used to calculate soma depth relative to drawings of the pia and/or white matter. Individual cells were also manually placed in the appropriate cortical region and layer within the Allen Mouse Common Coordinate Framework (CCF) by matching the 20x image of the slice with a “virtual” slice at an appropriate location and orientation within the CCF. Laminar locations (see Figure 2) were calculated by finding the path connecting pia and white matter that passed through the cell’s coordinate, identifying its distance to pia and white matter as well as position within its layer, then aligning those values to an average set of layer thicknesses. Using the DAPI image, laminar borders were also drawn for all reconstructed neurons.

#### Morphological reconstruction

Reconstructions of the dendrites and the full axon were generated for a subset of neurons with good quality transcriptomics, electrophysiology and biocytin fill. Reconstructions were generated based on a 3D image stack that was run through a Vaa3D-based image processing and reconstruction pipeline (Peng et al., 2010). Images were used to generate an automated reconstruction of the neuron using TReMAP (Zhou et al., 2016). Alternatively, reconstructions were created manually using the reconstruction software PyKNOSSOS (Ariadne-service) or the citizen neuroscience game Mozak (Mosak.science)(Roskams and Popović, 2016). Automated or manually-initiated reconstructions were then extensively manually corrected and curated using a range of tools (e.g., virtual finger, polyline) in the Mozak extension (Zoran Popovic, Center for Game Science, University of Washington) of Terafly tools (Peng et al., 2014; Bria et al., 2016) in Vaa3D. Every attempt was made to generate a completely connected neuronal structure while remaining faithful to image data. If axonal processes could not be traced back to the main structure of the neuron, they were left unconnected.

Before morphological feature analysis, reconstructed neuronal morphologies were expanded in the dimension perpendicular to the cut surface to correct for shrinkage (Deitcher et al., 2017; Egger et al., 2008) after processing. The amount of shrinkage was calculated by comparing the distance of the soma to the cut surface during recording and after fixation and reconstruction. A tilt angle correction was also performed based on the estimated difference (via CCF registration) between the slicing angle and the direct pia-white matter direction at the cell’s location (Gouwens et al., 2019).

#### Identifying transcriptomic types

For the reasons explained above, we used transcriptomes of dissociated cells from (Tasic et al., 2018) as reference dataset and mapped Patch-seq transcriptomes to the reference data to identify their cell types. The details are explained below and in Figure S1.

#### Preparation of reference cells and taxonomy tree

24,411 dissociated cells from VISp and ALM region and their corresponding cell types and a list of 4,020 most important genes were adopted from (Tasic et al., 2018). Only neuronal cells and their corresponding cell types from VISp region were selected from the above dataset (in total 13,464 cells and 93 cell types) as the reference dataset for mapping Patch-seq transcriptomes (Figure S1B). Two different mapping algorithms were used and are explained in the following.

#### Mapping on the reference taxonomy tree

The Patch-seq transcriptomes were mapped to the reference taxonomy tree in the top down manner, trying to resolve broad classes first, then the subtle subtypes (Figure S1C and D). For each individual cell, starting from the root and at each branch point, we computed its correlation with the reference cell types using the markers associated with the given branch point, and chose the most correlated branch. The process was repeated till reaching the leaves. To determine the confidence of mapping, we applied the 100 bootstrapping iterations at each branch point, and in each iteration, 70% of the reference cells and 70% of markers were randomly sampled for mapping. The percentage of times a cell was mapped to a given leaf or branch point in the reference taxonomy is defined as the corresponding mapping probability. For each cell, the mapping probability was sorted and the cell type that had highest probability was assigned as the corresponding cell type of that cell.

#### Mapping using a neural network classifier

A feedforward neural network classifier which receives gene expression data as input and predict the cell type for each individual cell, was built. The reference cells and their corresponding types were used to train the classifier. First the reference cells were divided into two groups of training (80%) and test (20%) sets. The input was the log_2_(CPM+1) gene expression data and the output was a probability vector which predict the type of each cell (see Figure S1E). After the model is trained, it was used to predict the cell type for the cells with Patch-seq recordings. The training on the dissociated cells and the subsequent prediction of the label of each cell (with Patch-seq recording) was done 100 time (each time the training was done on a new set of randomly selected 80% of dissociated cells). At the end, the percentage of times a cell was mapped to a given leaf or branch point in the reference taxonomy is defined as the corresponding mapping probability. For each cell, the mapping probability was sorted and the cell type that had highest probability was assigned as the corresponding cell type of that cell.

#### Assessing mapping quality

Due to presence of more significant contamination and dropout in the Patch-seq transcriptomes (see Figure S2), we occasionally observed ambiguous mapping of a cell to highly distinct cell types. To quantify the quality of mapping by considering what is “expected” and “unexpected” ambiguity of mapping between cell types we furthered categorized cells with Patch-seq recordings into three main groups: 1) Highly consistent cells, 2) Moderately consistent cells and 3) inconsistent cells. To do so, we first mapped the reference dissociated cells to the reference taxonomy to obtain the original or “expected” ambiguity which was present in the reference data set. By aggregating the mapping results and the clustering results (original cell types of the reference cells), a reference mapping probability matrix was constructed (Figure S1F). Then we examined whether the ambiguity in the mapping of Patch-seq transcriptomes is similar to this reference dissociated cells ambiguity by computing the Kullback-Leibler (KL) divergence (Berger et al., 2009) between the mapping probability distributions of cells with Patch-seq recordings and the reference mapping probability distribution. Cells with divergence greater than 2 were defined as “inconsistent” cells and removed from the analysis of this paper. If the KL divergence was less than 2, then the correlation of the gene expression between the cell and the reference cells from the same type was computed. If the correlation was less than 0.5, the cell was given an “inconsistent” label and was removed from the analysis. The quality of the remaining cells were further quantified by the probability of the mapping to one or more than one cell types. First the mapping probability of each individual cell to each of the 93 cell types were sorted. If the sum of the first two highest mapping probabilities was more than 70% and the ratio of the mapping probability to the first type with respect to the second type was more than 2, then the cell was labeled as “highly consistent” as it was very confidently mapped to the first type. Otherwise the cell was called a “moderately consistent” cell as it was still confidently mapped to one or more types (Figure S1G).

#### Jensen-Shannon divergence in transcriptomic features

The average, normalized CPM values for the ∼12000 most important genes for each cell-type were used to compute the Jensen-Shannon divergence (Lin, 1991) between each pair of cell types for dissociated cell and for cells with Patch-seq recordings. First, the CPM values of the selected genes was normalized to add up to 1 for each individual cell. Then, the mean CPM for each type was computed by averaging the CPM values over all the cells in each cell type and again normalizing to add up to 1 for each individual type. The normalized mean CPM values for each cell type was treated as the probability distribution of expression of the selected genes in each type. Then, the Jensen-Shannon divergence (*JSD*) was computed between each pair of cell types using their mean CPM values and the following equation

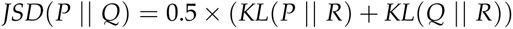

where *R* = 0.5 × (*P* + *Q*) denotes the midpoint of the probability vectors *P* and *Q*, and *KL*(*P R*) and *KL*(*Q R*) denote the Kullback-Leibler divergence of *R* from *P* and of *R* from *Q* (Lin, 1991; D.M. Endres, 2003).

### Electrophysiology feature analysis

Electrophysiological features were measured from responses elicited by short (3 ms) current pulses and long (1 s) current steps as previously described (Gouwens et al., 2019). Briefly, APs were detected by first identifying locations where the smoothed derivative of the membrane potential (dV/dt) exceeded 20 mV/ms, then refining based on several criteria including threshold-to-peak voltage and time differences and absolute peak height. After refinement, AP thresholds were re-calculated by finding the point for each AP where the dV/dt was 5% of the average maximal dV/dt across all APs. For each AP, threshold, peak, fast trough, and width (at half-height) were calculated, along with the upstroke/downstroke ratio (i.e., ratio of the peak upstroke dV/dt to the peak downstroke dV/dt).

Waveforms of the first APs elicited by the lowest-amplitude current pulses and steps were concatenated together for sparse principal component analysis (see below); the derivatives of these waveforms were also analyzed in this way. The voltage trajectory of the ISI was characterized by extracting the trace between the fast trough of the initial AP and the threshold of the following AP, normalizing their durations, and averaging together. AP features across the responses to long current steps were compared by binning APs into 20 ms intervals and averaging within a bin. If no APs fell within a bin, the value was interpolated from neighboring bins that had APs. This was done for stimulus amplitudes starting at a given cell’s rheobase, as well +40 pA and +80 pA above rheobase. If a sweep of an expected amplitude was unavailable (for example, if it failed one of the QC criteria), the missing values were interpolated from neighboring QC-passing sweeps. The instantaneous firing frequency (defined as the inverse of the ISI) was also binned and interpolated with 20 ms bins. The adaption of the instantaneous firing frequency was also analyzed by normalizing to the maximum rate observed during the step. Subthreshold responses to hyperpolarizing current steps were analyzed using downsampled (to 10 ms bins) membrane potential traces that were concatenated together. Responses from –10 pA to –90 pA steps (at a –40 pA interval) were used, and 200 ms of the time before and after the step were included as well. In addition, the largest amplitude hyperpolarizing step response was analyzed by normalizing to the minimum membrane potential reached and the baseline membrane potential.

Data sets were built by accumulating the feature vectors in each category (AP waveform, each AP feature across long steps, subthreshold response waveforms, etc). Sparse principal component analysis was performed separately on each data set. Principal components with an adjusted explained variance exceeding 1% were kept (typically 1 to 8 components from a given data set). Analysis of the data set yielded 44 total components. The components were z-scored and combined to form a reduced dimension feature matrix. The electrophysiological feature matrix used was visualized with a two-dimensional projection using Uniform Manifold Approximation and Projection (UMAP, (Becht et al., 2018))

### Morphology feature analysis

#### Calculating morphological features

Morphological features were calculated as previously described (Gouwens et al., 2019). Briefly, feature definitions were collected from prior studies (Scorcioni et al., 2008; Markram et al., 2015). Reconstructed neurons were aligned in the direction perpendicular to pia and white matter. Additional features, such as the laminar distribution of axon, were calculated from the aligned morphologies. While slice angle tilt and shrinkage correction were performed (see above), features predominantly determined by differences in the *z*-dimension were not analyzed to minimize technical artifacts due to *z*-compression of the slice after processing.

#### NBLAST scores

To assess the similarity of axonal morphologies directly, we used the NBLAST method of the nat.nblast R package (Costa et al., 2016; Kohl et al., 2013). This method compares a pair of morphologies by analyzing the distance and local direction of nearby points between the two structures. NBLAST similarity scores, which range from 0 (completely dissimilar) to 1 (completely identical), were calculated using version 1 of the algorithm with a standard deviation parameter of 10 microns. Morphologies were aligned on their somas in the *z*-direction (perpendicular to the cut surface of the slice) before NBLAST scoring. Scores were symmetrized by averaging the scores in each direction between a given pair. Representative morphologies for a t-type were chosen by selecting cells with high average NBLAST scores versus other cells of the same t-type.

### Feature correspondence analysis

#### Likelihood ratio analysis

The distributions of electrophysiological and morphological features between pairs of t-types were compared by a log-likelihood ratio method. The data from a pair of t-types were fit with either a one-component model (combining all cells) or a two-component model (separate for each t-type). The dimensionality of the data sets was reduced by performing principal components analysis (PCA) on the set of elecrophysiological or morphological features and keeping the top three principal components. Multivariate Gaussian fits with a diagonal covariance matrix were fit to these data sets, and the log-likelihood ratio (LLR) of the data under the two-component versus one-component model were calculated. Since the two-component model would produce a higher likelihood due to overfitting even if the underlying distributions were the same, the statistical significance of the observed LLR was assessed by parametric boostrap. Random samples of the same size as the actual data set were drawn 2,000 times from the one-component model, and LLRs were calculated in the same way as with the actual data. These bootstrapped LLRs were used to estimate the *p* values of the observed LLRs.

#### Supervised classification

Supervised classification was performed by training a random-forest classifier with 100 trees using balanced class weights, implemented with the scikit-learn Python package (Pedregosa et al., 2011). Out-of-bag confusion matrices and accuracies were reported.

#### Discriminant analysis

To assess the degree to which pairs of t-types exhibited discrete or continuous variation in electrophysiological and morphological feature space, we used a cross-validated discriminant analysis method based on one developed to address a similar question in transcriptomic feature space (Harris et al., 2018). For each pair of t-types, multivariate Guassians were fit to “training” cells from each t-type to describe a group of electrophysiological or morphological features (typically with one to six dimensions in each feature group). Then, the likelihood ratio was calculated for each held-out “test” cell between its own assigned t-type versus the other t-type. This produced a distribution of likelihood ratios for each t-type, and the separation of these distributions was summarized by a *d’* statistic, 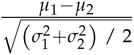, where *µ_i_* and *σ_i_* are the mean and standard deviations of the likelihood ratios for cells in t-type *i*. Larger values of *d’* indicate more separation between t-types.

#### ME-clustering

To identify types that had both consistent electrophysiological and morphological characteristics, we employed the same strategy developed for our previous work (Gouwens et al., 2019). Briefly, this method includes generating multiple combined electrophysiological and morphology feature matrices with different electrophysiological feature weights, then using multiple unsupervised clustering algorithms with each combined feature matrix (60 variations in total). A cell-wise co-clustering matrix was calculated from these variants, and consensus clusters were determined by iterative hierarchical clustering. Clusters were evaluated for stability by repeating the entire procedure on 90% subsamples and calculated the average Jaccard similarity; clusters with average Jaccard similarities below 0.5 were considered unstable and merged with another closest-matching cluster.

#### Spectral co-clustering

Spectral co-clustering was performed on the t-types to me-type adjacency matrix using the scikit-learn Python package (Pedregosa et al., 2011). This method simultaneously clustered the rows and columns of the matrix by treating it as a bipartite graph and approximating a normalized cut of the graph (Dhillon, 2001). This results in a block-diagonal structure, and the number of clusters was chosen to minimize the number of matrix elements that diverged from the block diagonal (i.e., the total number of non-zero elements outside the co-clusters and zero-valued elements within the co-clusters). Co-clustering was performed separately for each transcriptomic subclass.

#### Statistics and research design

No statistical methods were used to predetermine sample sizes, but the sample sizes here are similar to those reported in previous publications. No randomization was used during data collection as there was a single experimental condition for all acquired data. The different stimulus protocols were not presented in a randomized order. Data collection and analyses were not performed blind to the conditions of the experiments as there was a single experimental condition for all acquired data.

#### Data and software availability

Transcriptomic, electrophysiological, and morphological data supporting the findings of this study will be available online by mid-2020.

The custom electrophysiology data acquisition software (MIES) is available at https://github.com/alleninstitute/mies. The Vaa3D morphological reconstruction software, including the Mozak extension, is freely avail-able at http://www.vaa3d.org and its code is available at https://github.com/Vaa3D. The code for electrophysiological and morphological feature analysis and clustering is available as part of the opensource Allen SDK repository (https://github.com/AllenInstitute/AllenSDK), IPFX repository (https://github.com/alleninstitute/ipfx), and DRCME repository (https://github.com/alleninstitute/ drcme).

## Author Contributions

H.Z., C.K., and E.L. conceptualized the project; T.L.D. contributed to the generation and/or characterization of specific transgenic lines; L.E., M.R., and K.R. provided mouse colony management; E.B., T.C, K.C., N.D., L.E., M.K., J.S.,, and H.T. prepared brain slices; K.B., K.H., D.H., L.K., B.L, R.M,., L.N. A.O., R.R., and J.T. performed electrophysiological experiments; K.B., D.B, J.B., A.G., J.G., M.M., T.P., A.P., C.R., A.R., K.S., M.T., A.T, and K.W. prepared single cell RNASeq libraries; K.B., K.B, T.E., A.G., H.G., M.M., and D.P. processed slices for biocytin staining; S.D., N.D., R.E., M.G., M.H., K.N., L.P, and S.R. imaged biocytin-stained slices and cells; K.B., R.R, J.S., L.A., R.D., R.A.D., T.D., C.G., A.M.H., S.K., M.M., A.M., D.S., S.A.S., and G.W. reconstructed neurons and/or provided anatomical annotations; C.G., S.A.S., B.L., J.B., F.B., A.B., O.F., R.G., L.G., N.W.G., T.K.K., C.L., K.E.L., O.P., U.S., and Z.Y. performed analyses; S.A.S., B.L., J.B., L.G., T.K.K., O.P., A.O., N.D., and J.T. contributed to methods development studies; K.E.L., B.T., T.B., C.F., T.J., H.P., D.R., and Z.Z generated tools for pipeline data generation; L.E., and S.M.S. provided program management support; T.J., S.A.S., B.L., J.B., L.G., N.D., A.B., L.P., M.M., K.S., P.R.N., and H.Z. organized and managed pipeline data generation; A.B., K.S., F.B., N.W.G., D.F., L.N., A.S., and W.W. organized and managed pipeline data storage and processing; N.W.G., S.A.S., J.B., H.Z., J.T., A.A., M.J.H., C.K., E.L., and G.J.M. provided scientific direction; N.W.G., S.A.S., A.B., and F.B. prepared the figures; N.W.G., S.A.S., A.B., F.B., and G.J.M. wrote the manuscript in consultation with all others; H.Z., J.B., B.T., C.K., and U.S. provided substantial review and edits to the manuscript.

## Acknowledgments

We are grateful to Cathryn Cadwell, Federico Scala, and Andreas Tolias for sharing their Patch-seq experience with us in the early stage of our pipeline development. We appreciate feedback on the manuscript provided by Scott Owen, Jeremy Miller, and Brian Kalmbach. We thank Adrian Wanner for providing reconstruction services through Ariadne and Roy Szeto and Zoran Popovic for facilating the reconstruction work contributed by Mozak.science. Finally, we thank the Mozak citizen-scientists for their valuable contribution. The research was partially supported by several grant awards from institutes under the National Institutes of Health (NIH), including award numbers R01EY023173 from The National Eye Institute and U01MH105982 from the National Institute of Mental Health and Eunice Kennedy Shriver National Institute of Child Health & Human Development to H.Z. The content is solely the responsibility of the authors and does not necessarily represent the official views of NIH and its subsidiary institutes. This work was funded by the Allen Institute for Brain Science. We thank Sil Coulter, John Philips, and Allan Jones for leadership and guidance, and dedicate this paper to the vision, encouragement, and long-term support of our founder, Paul G. Allen.

## Supplementary Material

**Figure S1:**
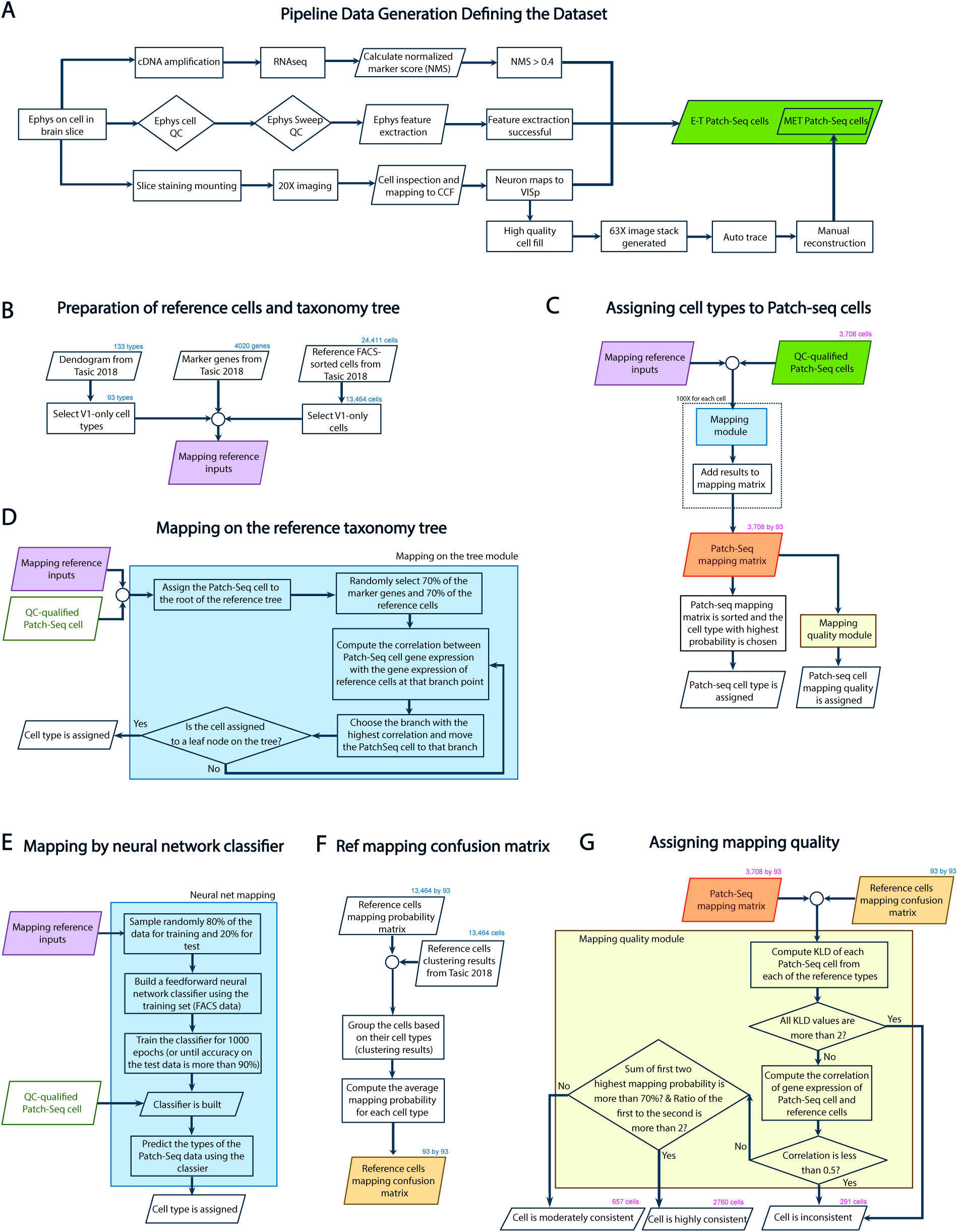
Data pipeline schematic. (A) Each cell underwent standardized collection of electrophysiology data, followed by a series of processing and quality control steps in parallel for each modality. Transcriptomic analysis included extraction and reverse transcription of the cell’s nuclear and cytosolic mRNA, followed by cDNA amplification, sequencing and evaluation of data quality based on normalized summed expression of “on”-type marker genes (NMS) adapted from the single-cell quality control measures in (Tripathy et al., 2018). Electrophysiological analysis included cell and sweep-level quality control gates, and automated extraction of feature vectors. Electrophysiology slices were stained, mounted onto coverslips, imaged at 20x and mapped to the Allen Mouse Common Coordinate Framework (CCF). QC-Qualified cells had an NMS *>* 0.4, passed electrophysiology quality control and feature extraction and had a confirmed soma location in primary visual cortex (VISp). A subset of cells with high-quality cell fills were further processed for morphological reconstruction. (B) Schematic of reference data set and mapping tree. (C) Assignment of t-types to cells assayed via Patch-seq recordings. QC-qualified cells from (A) were mapped on the reference dataset obtained in (B) using two different mapping algorithms (see (D) and (E)). (D) Schematic illustrating mapping procedure using the reference taxonomy tree. (E) Schematic illustrating mapping using a neural network classifier. (F) Calculation of reference mapping confusion matrix. The reference dissociated cells from (Tasic et al., 2018) were mapped to the reference taxonomy tree using the same method in (D), and the mapping probability matrix of those cells was computed. This probability matrix and the clustering results from (Tasic et al., 2018) of those dissociated cells were used as inputs to compute the reference confusion matrix. (G) Schematic illustrating the determination of mapping quality to each cell assayed via Patch-seq recordings.

**Figure S2:**
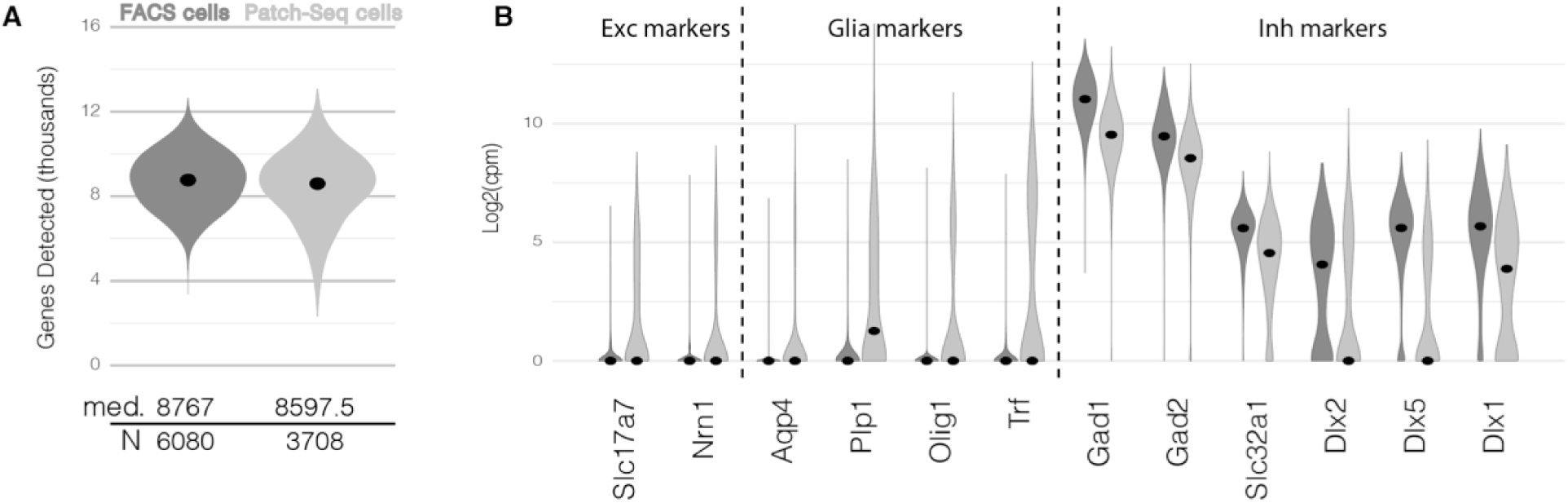
Comparing transcriptomic data quality of dissociated cells from (Tasic et al., 2018) and cells assayed via Patch-seq recordings. (A) Distribution of total number of genes detected in reference dissociated cells (dark grey) vs cells assayed via Patch-seq recordings (light grey) are presented by violin plots. The black dots are the medians. (B) Expression of selected excitatory marker genes (Slc17a7, Nrn1), Glia marker genes (Aqp4, Plp1, Olig1 and Trf) and inhibitory marker genes (Gad1, Gad2, Slc32a1, Dlx2, Dlx5 and Dlx1) are represented by violin plots.

**Figure S3:**
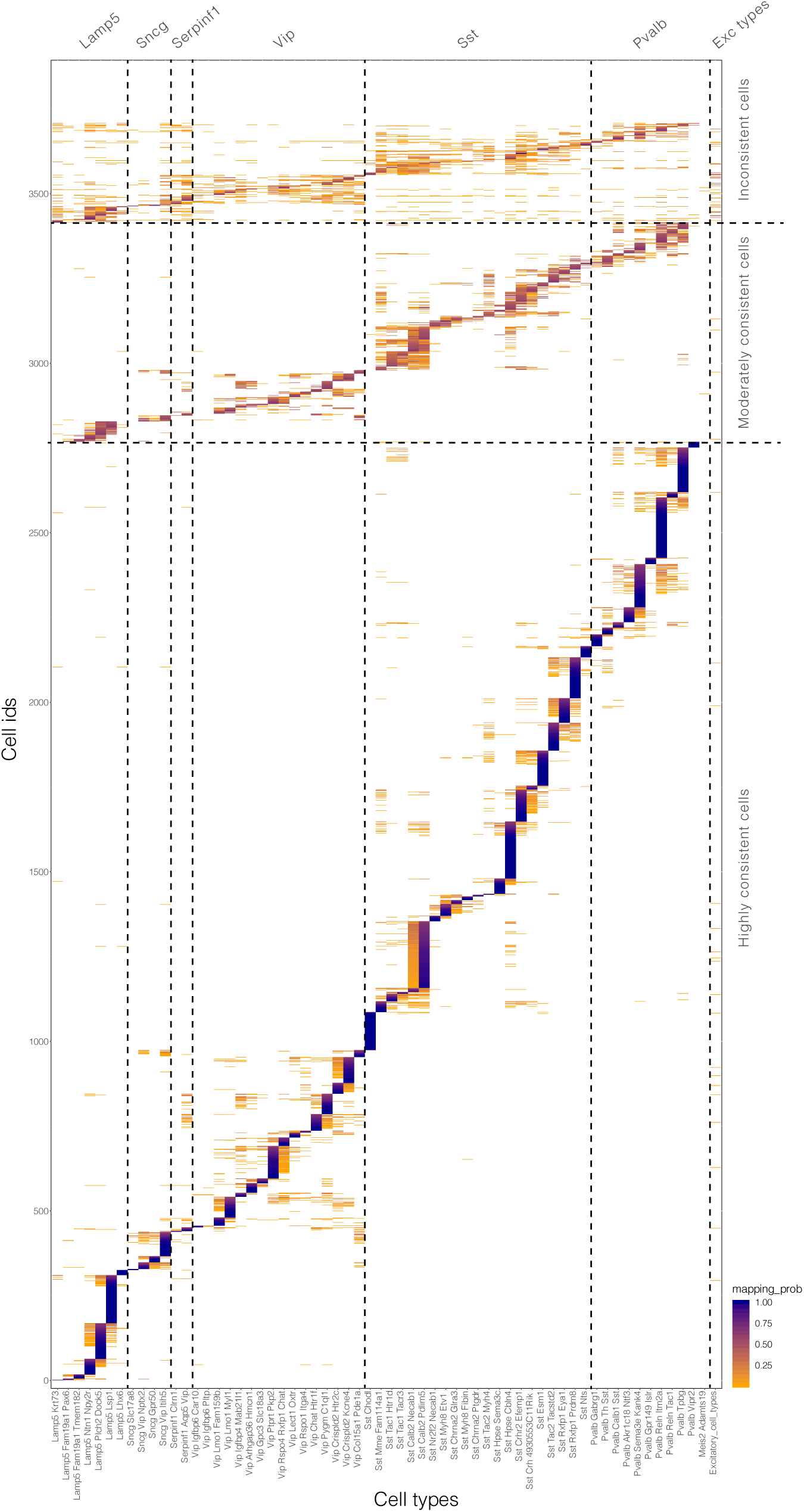
Mapping probability of Patch-seq transcriptomes on reference cell types from (Tasic et al., 2018). Mapping probability for three groups of cells described in Figure S1G (highly consistent, moderately consistent and inconsistent cells) using the reference tree mapping method (Figure S1D and Methods). Horizontal dashed lines divide mapping categories; vertical dashed lines divide transcriptomic subclasses.

**Figure S4:**
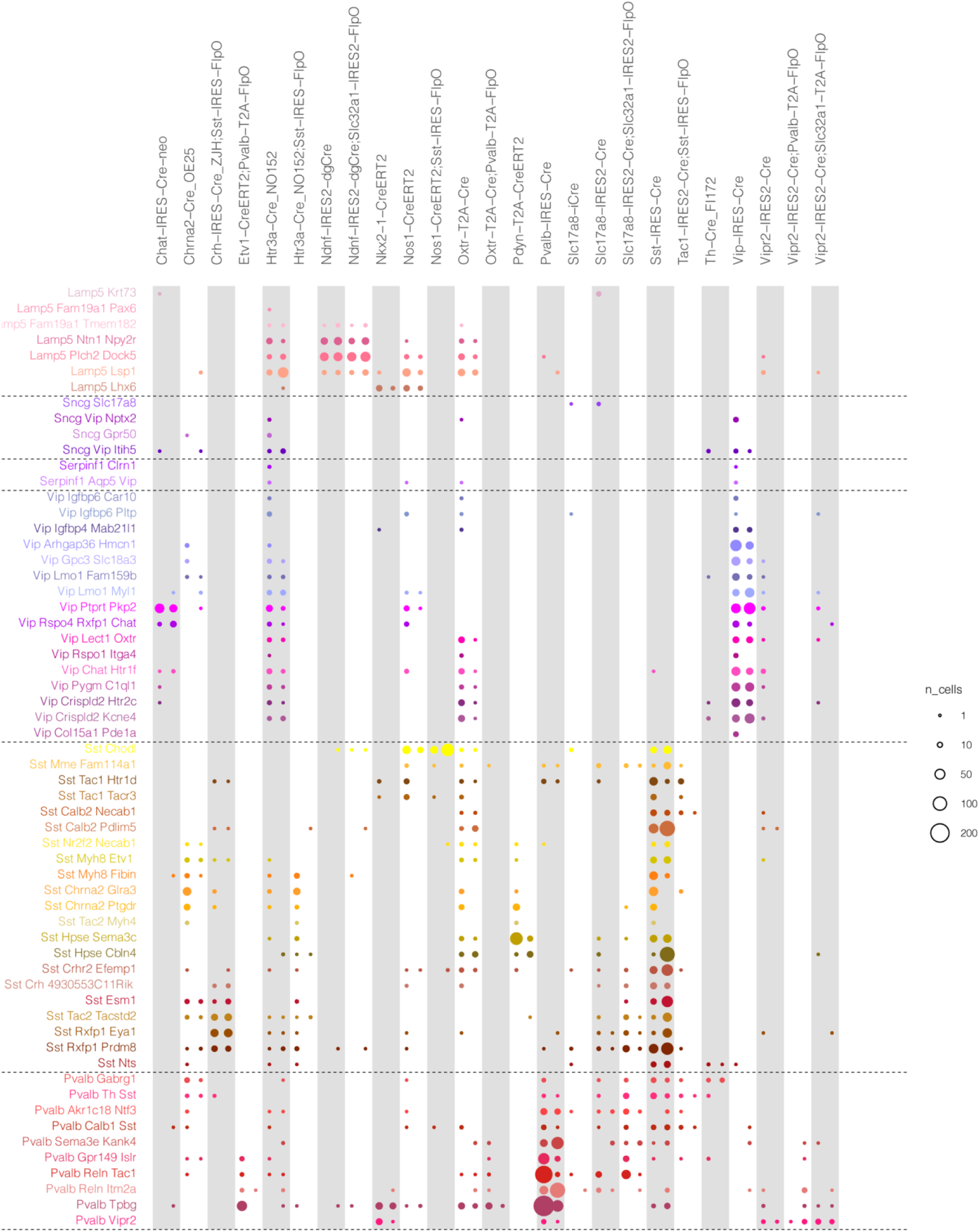
Common driver lines and their corresponding types in both data sets (cells assayed via Patch-seq recording and dissociated cells). Common driver line names that were present in this work and in the (Tasic et al., 2018) paper are listed on top (columns, n=26 driver lines) and cell types on the left (rows, n=60 t-types). Colored dots represent the numbers of cells detected for each type (cell numbers are proportional to dot area). Dissociated cells (n=2,607) on the left and cells assayed via Patchseq recordings (n=1,790 highly-consistent cells) are on the right in each column.

**Figure S5:**
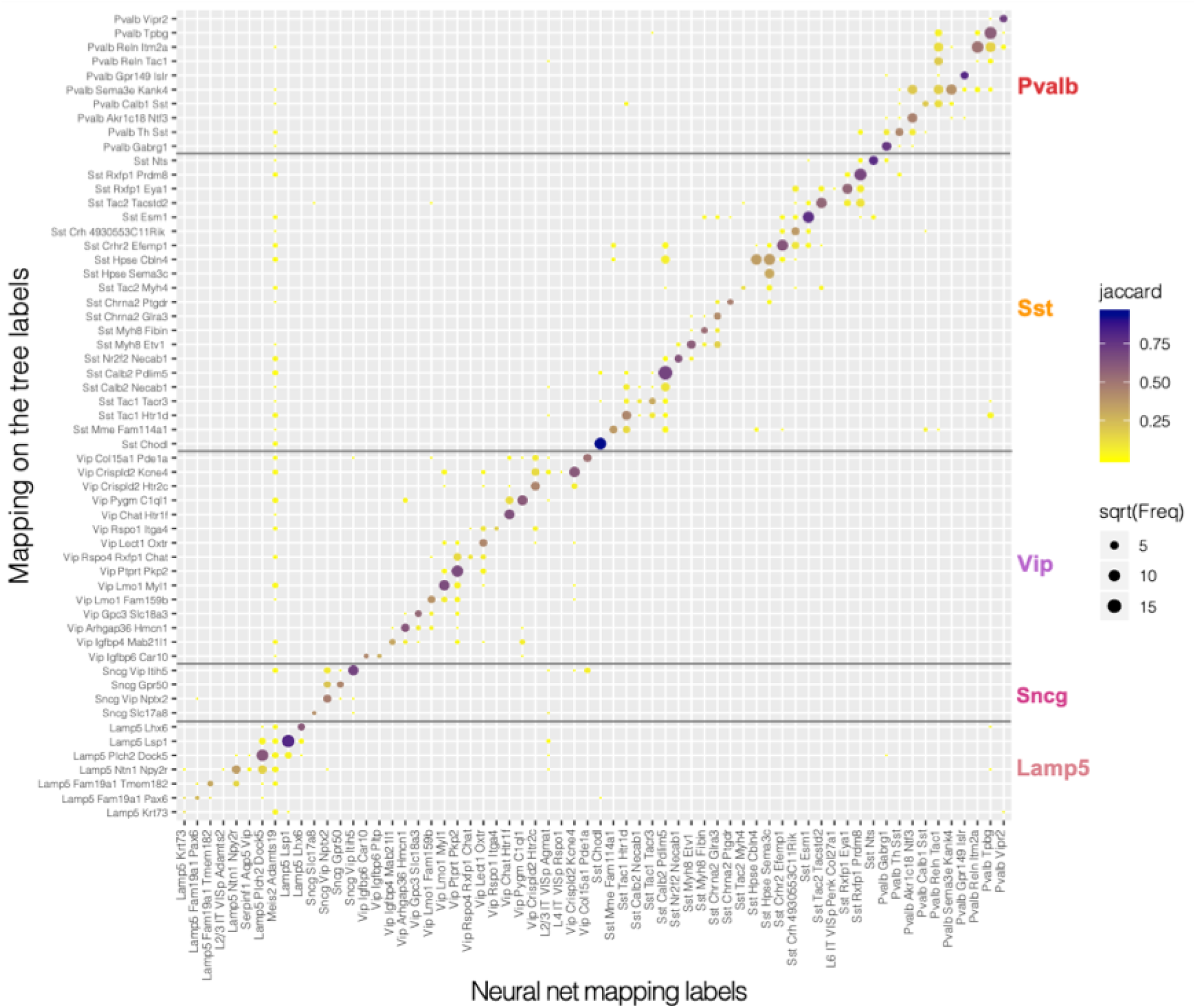
Comparison of mapping results using two independent algorithm. Result of the neural net method (see method and Figure S1) is shown on the x-axis and result of the mapping on tree method (see Methods and Figure S1) on the y-axis. We mapped all 3,708 cells from the current data set on to 93 cell types corresponding to the VISp region from previously published taxonomy (Tasic et al., 2018). The plot is showing the results for only the “highly consistent” cells (2,760 cells, see Figure S1G).

**Figure S6:**
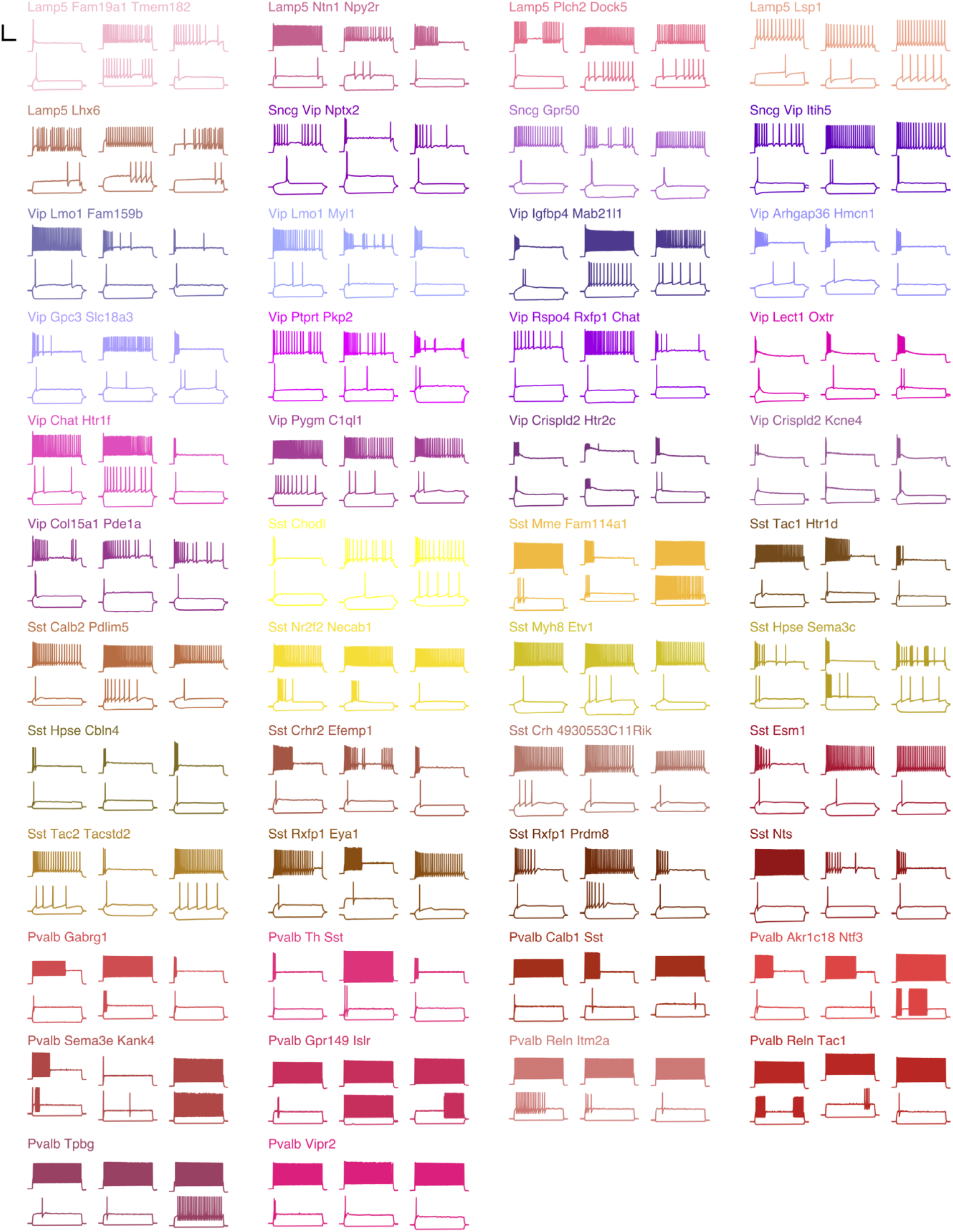
Example electrophysiological responses by t-type. Responses to 1 second-long current steps with stimulus amplitudes equal to −70 pA and rheobase for that cell (lower traces) and rheobase + 80 pA (upper trace). Three randomly-chosen examples shown for each t-type. Scale bar: vertical 50 mV, horizontal 250 ms.

**Figure S7:**
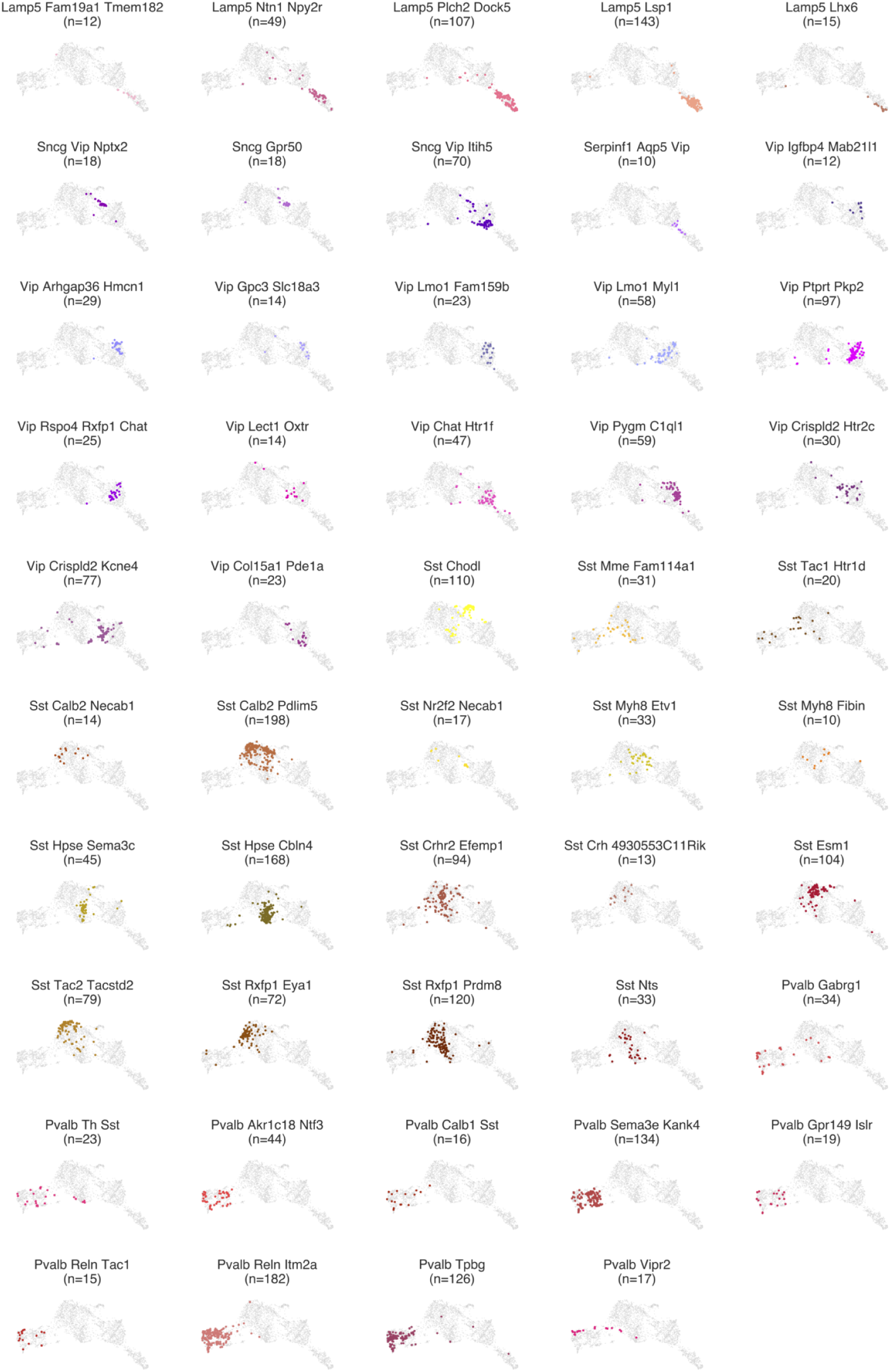
Electrophysiology UMAP plots by t-type. UMAP plots were generated using the 44 z-scored sparse principal component values collected from electrophysiology feature vectors. Each t-type with a minimum of 10 cells with highly-consistent mapping is shown (colors).

**Figure S8:**
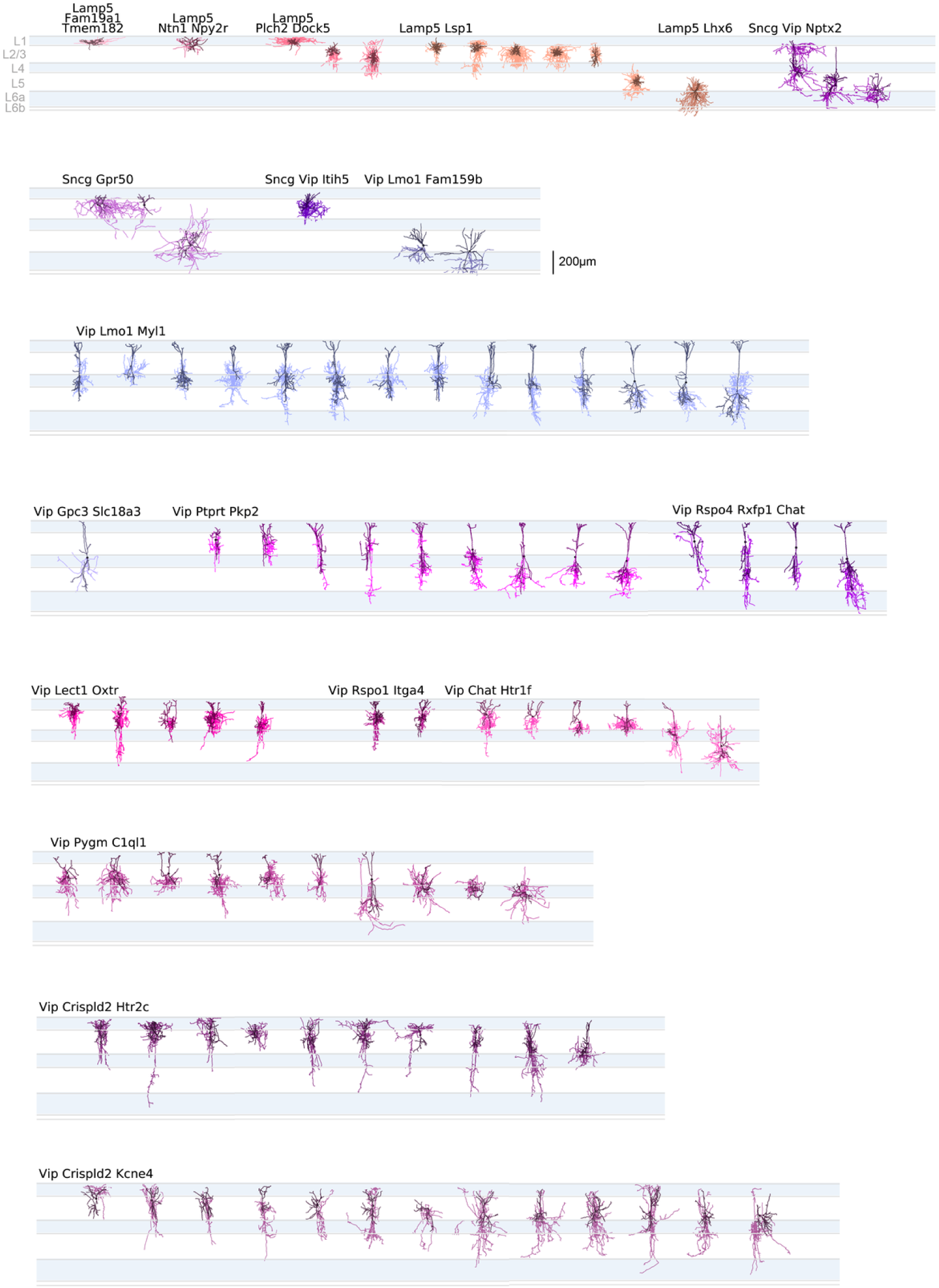
Lamp5 and Vip subclasses: morphological reconstructions ordered by t-type and relative soma depth. All morphological reconstructions from the Lamp5 and Vip subclasses aligned by layer to an average cortical template ordered by t-type and distance from pia within t-type. Dendrites are in darker colors, axon in lighter colors.

**Figure S9:**
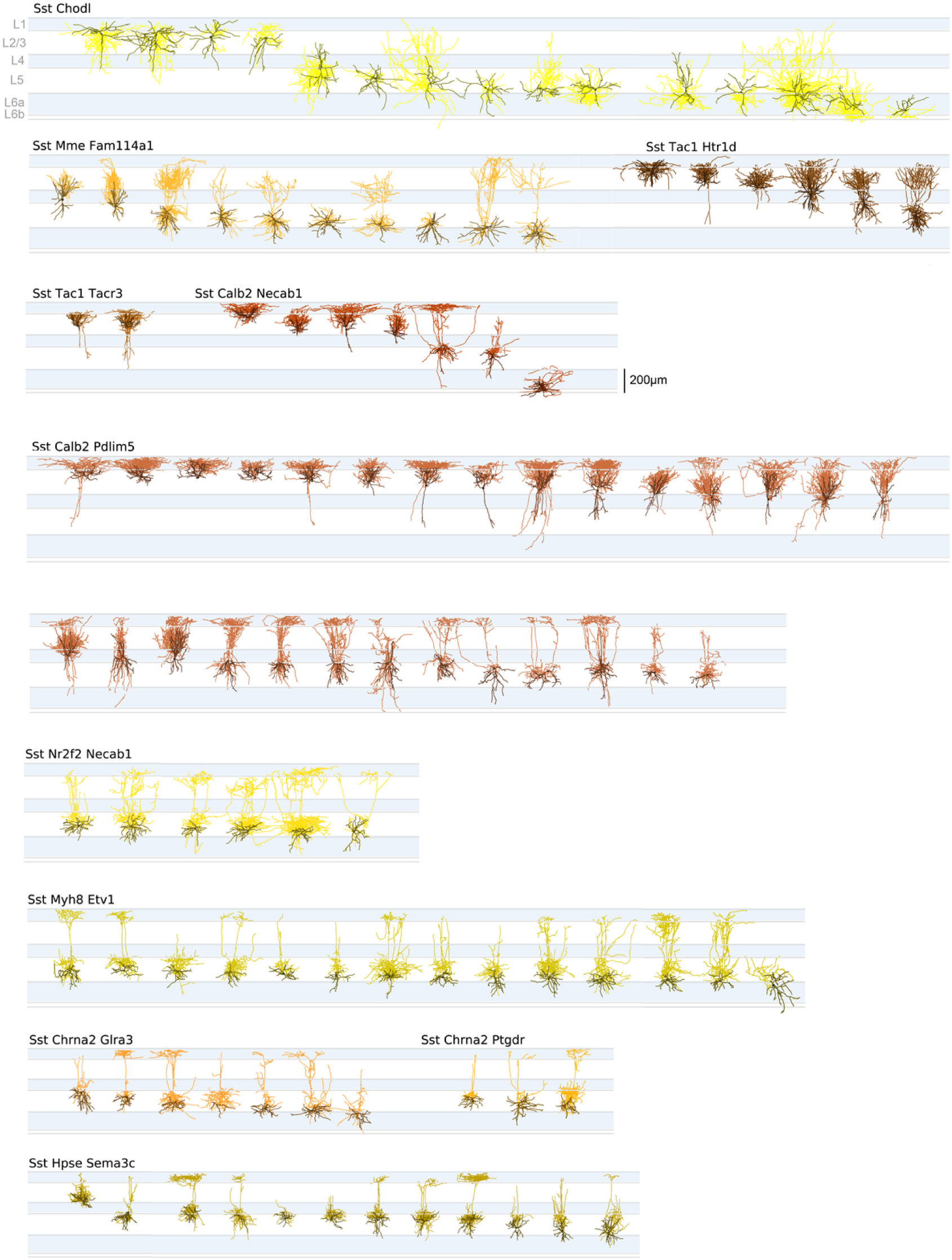
Sst subclass: morphological reconstructions ordered by t-type and relative soma depth. All morphological reconstructions from multiple t-types from the Sst subclass aligned by layer to an average cortical template. Dendrites are in darker colors, axon in lighter colors.

**Figure S10:**
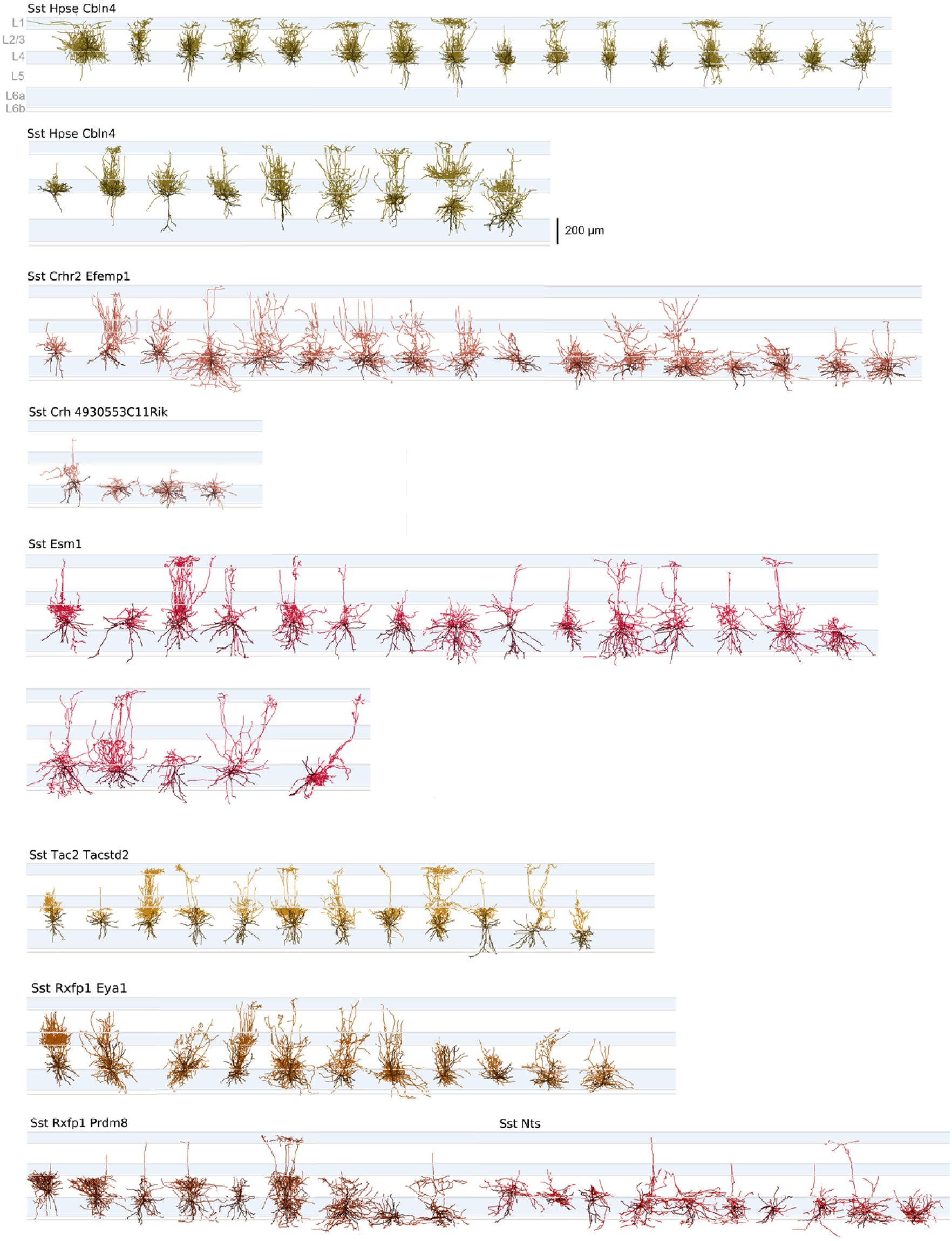
Sst subclass: morphological reconstructions ordered by t-type and relative soma depth. All morphological reconstructions from multiple t-types from the Sst subclass aligned by layer to an average cortical template. Dendrites are in darker colors, axon in lighter colors.

**Figure S11:**
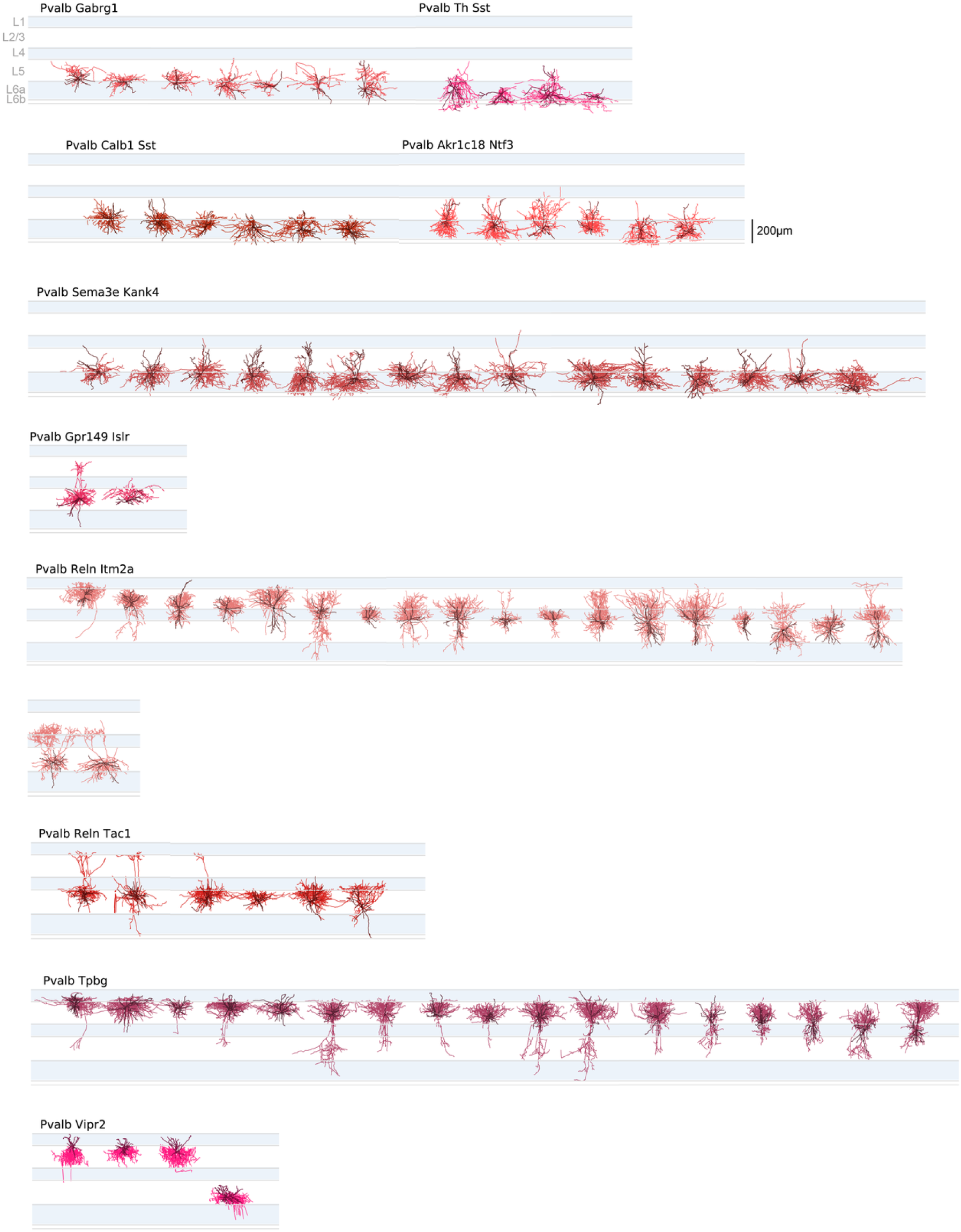
Pvalb subclass: morphological reconstructionsordered by t-type and relative soma depth. All morphological reconstructions from the Pvalb subclass aligned by layer to an average cortical template. Dendrites are in darker colors, axon in lighter colors.

**Figure S12:**
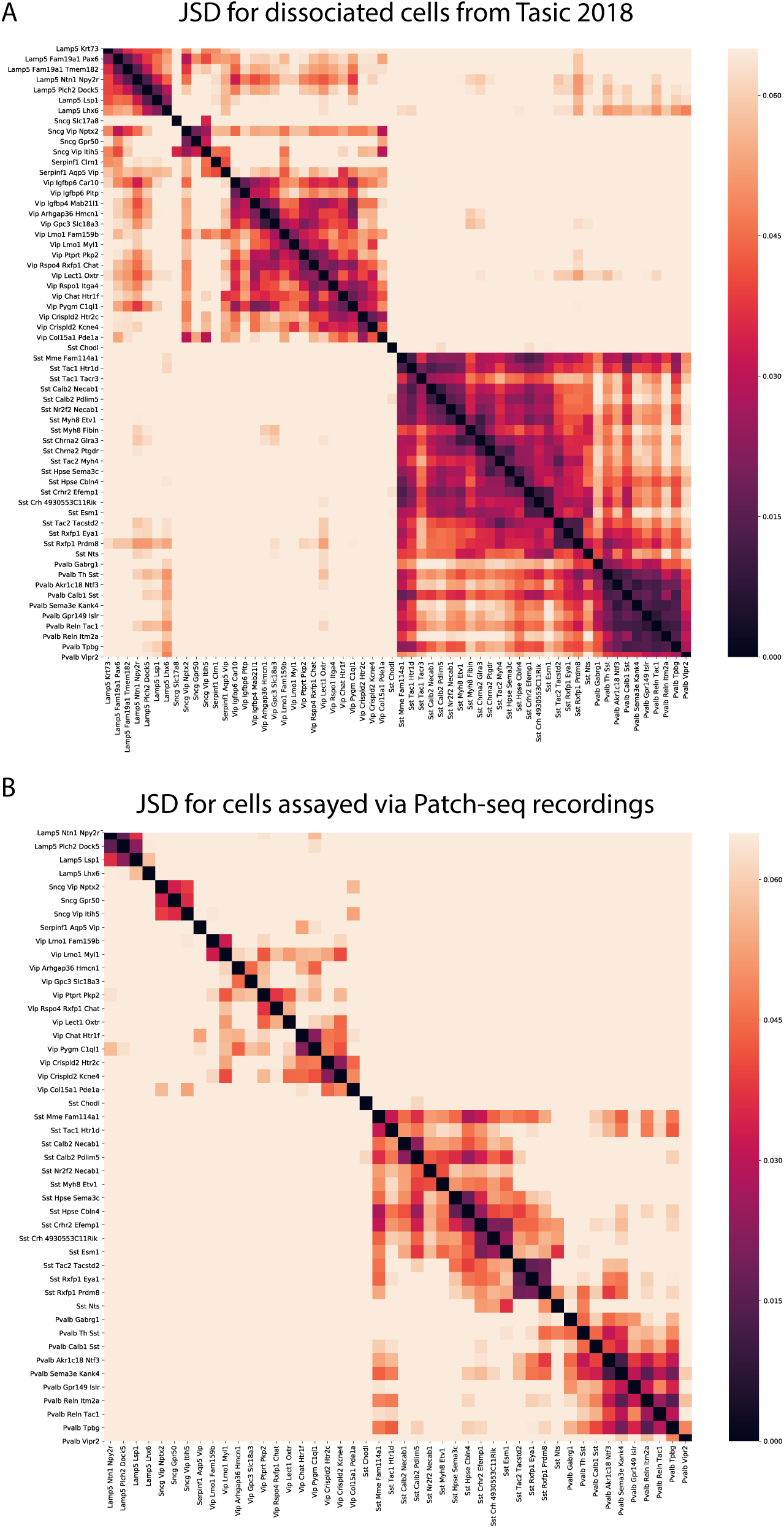
Jensen-Shannon Divergence (JSD). (A) JSD values between GABAergic cell types using the reference data set from (Tasic et al., 2018) (in total 6,080 GABAergic dissociated cells were used). (B) JSD values between Patch-seq GABAergic types. Only “highly consistent” cells and types with more than 10 cells are shown. All the JSD values more than 0.065 are shown as the same color as 0.065.

## References

1. Ascoli, G.A., Alonso-Nanclares, L., Anderson, S.A., Barrionuevo, G., Benavides-Piccione, R., Burkhalter, A., Buzsáki, G., Cauli, B., Defelipe, J., Fairén, A., et al. (2008). Petilla terminology: nomenclature of features of GABAergic interneurons of the cerebral cortex. Nat Rev Neurosci 9, 557–568.

2. Becht, E., McInnes, L., Healy, J., Dutertre, C.A., Kwok, I.W.H., Ng, L.G., Ginhoux, F., and Newell, E.W. (2018). Dimensionality reduction for visualizing single-cell data using UMAP. Nat Biotechnol 37, 38–44.

3. Berger, J.O., Bernardo, J.M., and Sun, D. (2009). The Formal Definition of Reference Priors. The Annals of Statistics 37, 905–938.

4. Bortone, D.S., Olsen, S.R., and Scanziani, M. (2014). Translaminar inhibitory cells recruited by layer 6 corticothalamic neurons suppress visual cortex. Neuron 82, 474–485.

5. Bria, A., Iannello, G., Onofri, L., and Peng, H. (2016). TeraFly: real-time three-dimensional visualization and annotation of terabytes of multidimensional volumetric images. Nat Methods 13, 192–194.

6. Cadwell, C.R., Palasantza, A., Jiang, X., Berens, P., Deng, Q., Yilmaz, M., Reimer, J., Shen, S., Bethge, M., Tolias, K.F., et al. (2016). Electrophysiological, transcriptomic and morphologic profiling of single neurons using Patch-seq. Nat Biotechnol 34, 199–203.

7. Cembrowski, M.S., and Spruston, N. (2019). Heterogeneity within classical cell types is the rule: lessons from hippocampal pyramidal neurons. Nat Rev Neurosci 20, 193–204.

8. Chen, K.H., Boettiger, A.N., Moffitt, J.R., Wang, S., and Zhuang, X. (2015). RNA imaging. Spatially resolved, highly multiplexed RNA profiling in single cells. Science 348, aaa6090.

9. Codeluppi, S., Borm, L.E., Zeisel, A., La, M.G., van, L.J.A., Svensson, C.I., and Linnarsson, S. (2018). Spatial organization of the somatosensory cortex revealed by osmFISH. Nat Methods 15, 932–935.

10. Coskun, A.F., and Cai, L. (2016). Dense transcript profiling in single cells by image correlation decoding. Nat Methods 13, 657–660.

11. Costa, M., Manton, J.D., Ostrovsky, A.D., Prohaska, S., and Jefferis, G.S. (2016). NBLAST: Rapid, Sensitive Comparison of Neuronal Structure and Construction of Neuron Family Databases. Neuron 91, 293–311.

12. D.M. Endres, J.E.S. (2003). A new metric for probability distributions. IEEE Trans. Inform. Theory 49.

13. DeFelipe, J., Ló pez-Cruz, P.L., Benavides-Piccione, R., Bielza, C., Larrañ aga, P., Anderson, S., Burkhalter, A., Cauli, B., Fairén, A., Feldmeyer, D., et al. (2013). New insights into the classification and nomenclature of cortical GABAergic interneurons. Nat Rev Neurosci 14, 202–216.

14. Deitcher, Y., Eyal, G., Kanari, L., Verhoog, M.B., Atenekeng, K.G.A., Mansvelder, H.D., de, K.C.P.J., and Segev, I. (2017). Comprehensive Morpho-Electrotonic Analysis Shows 2 Distinct Classes of L2 and L3 Pyramidal Neurons in Human Temporal Cortex. Cereb Cortex 27, 5398–5414.

15. Dhillon, I.S. (2001). Co-clustering documents and words using bipartite spectral graph partitioning. In Proceedings of the Seventh ACM SIGKDD International Conference on Knowledge Discovery and Data Mining KDD’01,(ACM Press),

16. Dobin, A., Davis, C.A., Schlesinger, F., Drenkow, J., Zaleski, C., Jha, S., Batut, P., Chaisson, M., and Gingeras, T.R. (2013). STAR: ultrafast universal RNA-seq aligner. Bioinformatics 29, 15–21.

17. Egger, V., Nevian, T., and Bruno, R.M. (2008). Subcolumnar dendritic and axonal organization of spiny stellate and star pyramid neurons within a barrel in rat somatosensory cortex. Cereb Cortex 18, 876–889.

18. Fuzik, J., Zeisel, A., Máté, Z., Calvigioni, D., Yanagawa, Y., Szabó, G., Linnarsson, S., and Harkany, T. (2016). Integration of electrophysiological recordings with single-cell RNA-seq data identifies neuronal subtypes. Nat Biotechnol 34, 175–183.

19. Fö ldy, C., Darmanis, S., Aoto, J., Malenka, R.C., Quake, S.R., and Sü dhof, T.C. (2016). Single-cell RNAseq reveals cell adhesion molecule profiles in electrophysiologically defined neurons. Proc Natl Acad Sci U S A 113, E5222–31.

20. Gouwens, N.W., Sorensen, S.A., Berg, J., Lee, C., Jarsky, T., Ting, J., Sunkin, S.M., Feng, D., Anastassiou, C.A., Barkan, E., et al. (2019). Classification of electrophysiological and morphological neuron types in the mouse visual cortex. Nat Neurosci 22, 1182–1195.

21. Harris, K.D., Hochgerner, H., Skene, N.G., Magno, L., Katona, L., Bengtsson, G.C., Somogyi, P., Kessaris, N., Linnarsson, S., and Hjerling-Leffler, J. (2018). Classes and continua of hippocampal CA1 inhibitory neurons revealed by single-cell transcriptomics. PLoS Biol 16, e2006387.

22. He, M., Tucciarone, J., Lee, S., Nigro, M.J., Kim, Y., Levine, J.M., Kelly, S.M., Krugikov, I., Wu, P., Chen, Y., et al. (2016). Strategies and Tools for Combinatorial Targeting of GABAergic Neurons in Mouse Cerebral Cortex. Neuron 92, 555.

23. Hodge, R.D., Bakken, T.E., Miller, J.A., Smith, K.A., Barkan, E.R., Graybuck, L.T., Close, J.L., Long, B., Johansen, N., Penn, O., et al. (2019). Conserved cell types with divergent features in human versus mouse cortex. Nature 573, 61–68.

24. Huang, Z.J., and Paul, A. (2019). The diversity of GABAergic neurons and neural communication elements. Nat Rev Neurosci 20, 563–572.

25. Jiang, X., Shen, S., Cadwell, C.R., Berens, P., Sinz, F., Ecker, A.S., Patel, S., and Tolias, A.S. (2015). Principles of connectivity among morphologically defined cell types in adult neocortex. Science 350, aac9462–aac9462.

26. Kawaguchi, Y., and Kubota, Y. (1996). Physiological and morphological identification of somatostatinor vasoactive intestinal polypeptide-containing cells among GABAergic cell subtypes in rat frontal cortex. J Neurosci 16, 2701–2715.

27. Kebschull, J.M., Garcia, da S.P., Reid, A.P., Peikon, I.D., Albeanu, D.F., and Zador, A.M. (2016). HighThroughput Mapping of Single-Neuron Projections by Sequencing of Barcoded RNA. Neuron 91, 975–987.

28. Klausberger, T., and Somogyi, P. (2008). Neuronal diversity and temporal dynamics: the unity of hippocampal circuit operations. Science 321, 53–57.

29. Kohl, J., Ostrovsky, A.D., Frechter, S., and Jefferis, G.S. (2013). A bidirectional circuit switch reroutes pheromone signals in male and female brains. Cell 155, 1610–1623.

30. Kuan, L., Li, Y., Lau, C., Feng, D., Bernard, A., Sunkin, S.M., Zeng, H., Dang, C., Hawrylycz, M., and Ng, L. (2015). Neuroinformatics of the Allen Mouse Brain Connectivity Atlas. Methods 73, 4–17.

31. Lawrence, M., Huber, W., Pagès, H., Aboyoun, P., Carlson, M., Gentleman, R., Morgan, M.T., and Carey, V.J. (2013). Software for computing and annotating genomic ranges. PLoS Comput Biol 9, e1003118.

32. Lee, J.H., Daugharthy, E.R., Scheiman, J., Kalhor, R., Yang, J.L., Ferrante, T.C., Terry, R., Jeanty, S.S., Li, C., Amamoto, R., et al. (2014). Highly multiplexed subcellular RNA sequencing in situ. Science 343, 1360–1363.

33. Lin, J. (1991). Divergence Measures Based on the Shannon Entropy. IEEE Trans. Inform. Theory 37.

34. Markram, H., Muller, E., Ramaswamy, S., Reimann, M.W., Abdellah, M., Sanchez, C.A., Ailamaki, A., Alonso-Nanclares, L., Antille, N., Arsever, S., et al. (2015). Reconstruction and Simulation of Neocortical Microcircuitry. Cell 163, 456–492.

35. McGarry, L.M., Packer, A.M., Fino, E., Nikolenko, V., Sippy, T., and Yuste, R. (2010). Quantitative classification of somatostatin-positive neocortical interneurons identifies three interneuron subtypes. Front Neural Circuits 4, 12.

36. Moffitt, J.R., Bambah-Mukku, D., Eichhorn, S.W., Vaughn, E., Shekhar, K., Perez, J.D., Rubinstein, N.D., Hao, J., Regev, A., Dulac, C., et al. (2018). Molecular, spatial, and functional single-cell profiling of the hypothalamic preoptic region. Science 362.

37. Muñ oz, W., Tremblay, R., Levenstein, D., and Rudy, B. (2017). Layer-specific modulation of neocortical dendritic inhibition during active wakefulness. Science 355, 954–959.

38. Naka, A., Veit, J., Shababo, B., Chance, R.K., Risso, D., Stafford, D., Snyder, B., Egladyous, A., Chu, D., Sridharan, S., et al. (2019). Complementary networks of cortical somatostatin interneurons enforce layer specific control. Elife 8.

39. Neher, E. (1992). Correction for liquid junction potentials in patch clamp experiments. Methods Enzymol 207, 123–131.

40. Nigro, M.J., Hashikawa-Yamasaki, Y., and Rudy, B. (2018). Diversity and Connectivity of Layer 5 Somatostatin-Expressing Interneurons in the Mouse Barrel Cortex. J Neurosci 38, 1622–1633.

41. Paul, A., Crow, M., Raudales, R., He, M., Gillis, J., and Huang, Z.J. (2017). Transcriptional Architecture of Synaptic Communication Delineates GABAergic Neuron Identity. Cell 171, 522–539.e20.

42. Pedregosa, F., Varoquaux, G., Gramfort, A., Michel, V., Thirion, B., Grisel, O., Blondel, M., Prettenhofer, P., Weiss, R., Dubourg, V., et al. (2011). Scikit-learn: Machine Learning in Python. Journal of Machine Learning Research 12, 2825–2830.

43. Peng, H., Bria, A., Zhou, Z., Iannello, G., and Long, F. (2014). Extensible visualization and analysis for multidimensional images using Vaa3D. Nat Protoc 9, 193–208.

44. Peng, H., Ruan, Z., Long, F., Simpson, J.H., and Myers, E.W. (2010). V3D enables real-time 3D visualization and quantitative analysis of large-scale biological image data sets. Nat Biotechnol 28, 348–353.

45. Prö nneke, A., Scheuer, B., Wagener, R.J., Mö ck, M., Witte, M., and Staiger, J.F. (2015). Characterizing VIP Neurons in the Barrel Cortex of VIPcre/tdTomato Mice Reveals Layer-Specific Differences. Cereb Cortex 25, 4854–4868.

46. Que, L., Lukacsovich, D., and Fö ldy, C. (2020). Transcriptomic homogeneity and an age-dependent onset of hemoglobin expression characterize morphological PV types.

47. Roskams, J., and Popović, Z. (2016). Power to the People: Addressing Big Data Challenges in Neuroscience by Creating a New Cadre of Citizen Neuroscientists. Neuron 92, 658–664.

48. Saunders, A., Macosko, E.Z., Wysoker, A., Goldman, M., Krienen, F.M., de, R.H., Bien, E., Baum, M., Bortolin, L., Wang, S., et al. (2018). Molecular Diversity and Specializations among the Cells of the Adult Mouse Brain. Cell 174, 1015–1030.e16.

49. Scala, F., Kobak, D., Shan, S., Bernaerts, Y., Laturnus, S., Cadwell, C.R., Hartmanis, L., Froudarakis, E., Castro, J.R., Tan, Z.H., et al. (2019). Layer 4 of mouse neocortex differs in cell types and circuit organization between sensory areas. Nat Commun 10, 4174.

50. Scorcioni, R., Polavaram, S., and Ascoli, G.A. (2008). L-Measure: a web-accessible tool for the analysis, comparison and search of digital reconstructions of neuronal morphologies. Nat Protoc 3, 866–876.

51. Shekhar, K., Lapan, S.W., Whitney, I.E., Tran, N.M., Macosko, E.Z., Kowalczyk, M., Adiconis, X., Levin, J.Z., Nemesh, J., Goldman, M., et al. (2016). Comprehensive Classification of Retinal Bipolar Neurons by Single-Cell Transcriptomics. Cell 166, 1308–1323.e30.

52. Tasic, B., Menon, V., Nguyen, T.N., Kim, T.K., Jarsky, T., Yao, Z., Levi, B., Gray, L.T., Sorensen, S.A., Dolbeare, T., et al. (2016). Adult mouse cortical cell taxonomy revealed by single cell transcriptomics. Nature Neuroscience 19, 335–346.

53. Tasic, B., Yao, Z., Graybuck, L.T., Smith, K.A., Nguyen, T.N., Bertagnolli, D., Goldy, J., Garren, E., Economo, M.N., Viswanathan, S., et al. (2018). Shared and distinct transcriptomic cell types across neocortical areas. Nature 563, 72–78.

54. Tebaykin, D., Tripathy, S.J., Binnion, N., Li, B., Gerkin, R.C., and Pavlidis, P. (2018). Modeling sources of interlaboratory variability in electrophysiological properties of mammalian neurons. J Neurophysiol 119, 1329–1339.

55. Tremblay, R., Lee, S., and Rudy, B. (2016). GABAergic Interneurons in the Neocortex: From Cellular Properties to Circuits. Neuron 91, 260–292.

56. Tripathy, S.J., Toker, L., Bomkamp, C., Mancarci, B.O., Belmadani, M., and Pavlidis, P. (2018). Assessing Transcriptome Quality in Patch-Seq Datasets. Front Mol Neurosci 11, 363.

57. Wang, Y., Toledo-Rodriguez, M., Gupta, A., Wu, C., Silberberg, G., Luo, J., and Markram, H. (2004). Anatomical, physiological and molecular properties of Martinotti cells in the somatosensory cortex of the juvenile rat. J Physiol 561, 65–90.

58. Zeisel, A., Hochgerner, H., Lö nnerberg, P., Johnsson, A., Memic, F., Zwan, J. van der, Häring, M., Braun, E., Borm, L.E., Manno, G.L., et al. (2018). Molecular Architecture of the Mouse Nervous System. Cell 174, 999–1014.e22.

59. Zeisel, A., Munoz-Manchado, A.B., Codeluppi, S., Lonnerberg, P., Manno, G.L., Jureus, A., Marques, S., Munguba, H., He, L., Betsholtz, C., et al. (2015). Cell types in the mouse cortex and hippocampus revealed by single-cell RNA-seq. Science 347, 1138–1142.

60. Zeng, H., and Sanes, J.R. (2017). Neuronal cell-type classification: challenges, opportunities and the path forward. Nat Rev Neurosci 18, 530–546.

61. Zhou, X., Mansori, I., Fischer, T., Witte, M., and Staiger, J.F. (2020). Characterizing the morphology of somatostatin-expressing interneurons and their synaptic innervation pattern in the barrel cortex of the GFP-expressing inhibitory neurons mouse. J Comp Neurol 528, 244–260.

62. Zhou, Z., Liu, X., Long, B., and Peng, H. (2016). TReMAP: Automatic 3D Neuron Reconstruction Based on Tracing, Reverse Mapping and Assembling of 2D Projections. Neuroinformatics 14, 41–50.

